# Whole-brain mapping of effective connectivity by fMRI with cortex-wide patterned optogenetics

**DOI:** 10.1101/2022.07.12.499420

**Authors:** Seonghoon Kim, Hyun Seok Moon, Thanh Tan Vo, Chang-Ho Kim, Geun Ho Im, Myunghwan Choi, Seong-Gi Kim

## Abstract

Functional magnetic resonance imaging (fMRI) with optogenetic neural manipulation is a powerful tool that enables brain-wide mapping of effective functional networks. To achieve flexible manipulation of neural excitation throughout the mouse cortex, we incorporated spatiotemporal programmable optogenetic stimuli generated by a digital micromirror device into an MR scanner via an optical fiber bundle for the first time. This approach offered versatility in space and time in planning the photostimulation pattern, combined with *in situ* optical imaging and cell-type or circuit-specific genetic targeting in individual mice. Brain-wide effective connectivity obtained by fMRI with optogenetic stimulation of atlas-based cortical regions is generally congruent with anatomically defined axonal tracing data but is affected by the types of anesthetics that act selectively on specific connections. fMRI combined with flexible optogenetics opens a new path to investigate dynamic changes in functional brain states in the same animal through high-throughput brain-wide effective connectivity mapping.

## INTRODUCTION

Noninvasive mapping of functional architecture is critical for investigating causally important changes in brain circuitry associated with behavioral and pathological changes. For this purpose, hemodynamic-based functional magnetic resonance imaging (fMRI) has been widely used as a surrogate of neural activity for noninvasive mapping of functional connectivity (FC) by the degree of synchrony among fMRI time series among anatomically distinct brain regions^1,2^. FC is similar in humans between task and rest conditions and can be used as a fingerprint that predicts cognitive behavior in individual subjects^3,4^. However, to better understand FC at the circuit level, effective connectivity (EC), implying causal influence of one brain region on another, is required^5^. However, the limited temporal resolution of fMRI (typically ~1 s) and slow hemodynamic function make it difficult to infer the direction of influence. To overcome this limitation, the modulation of neuronal activity in well-defined regions can be used to trigger downstream fMRI responses, allowing us to determine the direction and strength of EC.

Optogenetics, a means of using light to control the activity of genetically specified neurons^6^, has been combined with whole-brain fMRI or cortex-wide optical intrinsic signal (OIS) imaging to obtain EC data^7–12^. In optogenetic fMRI (ofMRI), light delivery has been implemented by using pigtailed optical fibers surgically implanted in the target brain region. Although this approach is beneficial for targeting deep brain areas, the implantation of the optical fiber severely restricts the flexibility of experimental designs; the location and distribution of optogenetic stimuli are largely predefined during surgical implantation. Moreover, the restricted number of surgically implantable optical fibers limits the experimental throughput, typically, to one target site per animal. Although several approaches have been proposed to achieve more flexible spatiotemporal light delivery for optogenetics, such as focal beam scanning^13,14^ or structured illumination^15–18^, these approaches have not yet been combined with fMRI due to the difficulty in the implementation of MR-compatible and adjustable optics within the narrow MR bore.

Here, we developed a novel opto-MR hybrid system integrating widefield cortex-wide optical imaging and patterned optogenetics with whole-brain fMRI for rapid mapping of EC in cell type-specific channelrhodopsin-2 (ChR2) transgenic mice. Spatiotemporally flexible light patterns were generated with a programmable digital micromirror device (DMD) and delivered through a high-density optical fiber bundle with 100k cores to the cortex of a mouse placed in an MR scanner. The optical system achieved a spatial resolution of ~40 μm with coverage of the whole mouse cranium for targeted optogenetics. We were able to obtain a set of whole-brain ECs from nine atlas-based cortical regions of a single Thy1-ChR2 mouse in a single experimental session. Optogenetic fMRI-based EC closely resembles the monosynaptic axonal projection patterns demonstrated in the Allen Mouse Connectivity Atlas, although they are modulated by the effects of anesthesia. We further examined functionally distinct ECs under the altered brain state induced by different anesthetics. Our approach demonstrates an ability to investigate brain state-dependent brain-wide EC changes in live animals.

## RESULTS

### System development for patterned ofMRI

For comprehensive EC mapping, we developed a hybrid system that allows spatiotemporally flexible cortex-wide optogenetics in conjunction with whole-brain fMRI readout (**Fig. 1a, Extended Data Fig. 1**). To avoid the influence of the magnetic field on optoelectronics, we designed an optic system consisting of two spatially separated modules: i) a patterned optogenetics module that was placed outside of the MR bore and ii) a projection module placed within the MR bore. The patterned optogenetics module generated an arbitrary light field using a DMD composed of ~4 million individually programmable micromirrors (2560×1600 pixels) and incorporated widefield optical imaging, thus allowing visualization of the cortical surface in either reflectance or fluorescence mode for *in situ* cortex-wide readout of structural or functional optical contrasts. The projection module comprises a projection lens, a right-angle prism mirror, and a 3-axis translational stage made of an MR-compatible plastic (polyether ether ketone, PEEK). The module is attached to the vendor-supplied MR animal cradle. A fiber bundle containing 100k cores (length: 3 m, diameter: 1.4 mm) optically connects the two modules by relaying the light fields for imaging and optogenetics bidirectionally. We confirmed that the projection module did not introduce any noticeable artifacts in single-shot gradient-echo echo-planar images (EPIs) or T2-weighted MR images (**Supplementary Fig. 1**).

**Fig. 1.**
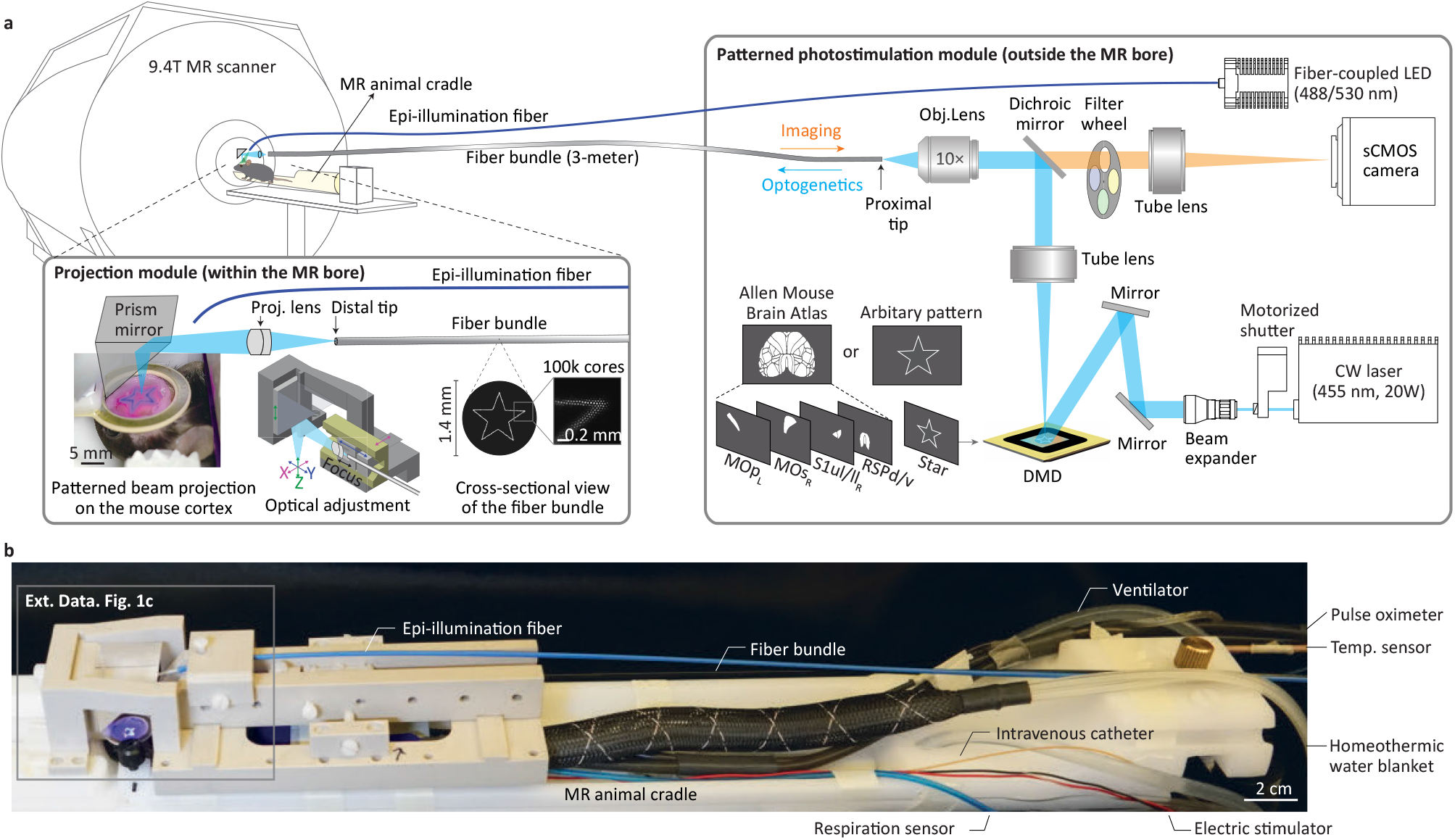
Overall design of the patterned optogenetic fMRI. **a**, Schematic illustration of the overall patterned photostimulation system combined with the fMRI setup. A patterned photostimulation module consisting of optical imaging and optogenetic beam paths is connected via a fiber bundle to a projection module attached to the MR animal cradle. The optogenetic beam emitted from a laser is expanded and relayed to a DMD. The beam is spatially sculpted by computer-generated binary patterns on the DMD and relayed to an objective lens (10x, NA 0.3) by passing through a tube lens (TL) and a dichroic mirror (DM). The arbitrary- or cortex regionshaped beam patterns are projected onto the proximal end of a 3-meter-long fiber bundle. The beam patterns emitted from the distal end are projected on the cortex of the thinned-skull mouse brain placed under the projection module. Reflectance or fluorescence imaging modes can be modified by changing the bandpass filter (BF) on a filter wheel and the color of a fiber-coupled LED. **b**, Photograph of the projection module attached to an MR animal cradle. Animal life-support equipment (pulse oximeter, temperature sensor, respiration sensor, intravenous catheter, and homeothermic water blanket) and an electric stimulator were connected to the cradle. The surface coil was placed on the head of the mouse around the thinned-skull window. An expanded view of the projection module is available in **Extended Data Fig. 1c**.

We next characterized the developed optic system (**Extended Data Fig. 2**). On the sample plane, the field of view was ~14 mm in diameter, and the depth of field was ~2.76 mm; these parameters were sufficient to cover the whole mouse cranium, which was ~10 mm in size and had a maximum depth difference of 1.1–1.3 mm^19^. The spatial resolution for imaging and optogenetics was ~40 μm, which was limited by the core pitch of the fiber bundle (~4 μm) multiplied by the magnification of the projection module (10x). Optical irradiance for patterned optogenetics reached up to 32 mW/mm^2^, which was sufficient to achieve physiological irradiance for optogenetics in deep cortical layers. The DMD supported a refresh rate of up to 9.5 kHz, providing submillisecond temporal precision.

For the animal model, we prepared a cortex-wide thinned-skull window, either in a transgenic or a viral-mediated ChR2-expressing mouse, and further introduced several modifications to minimize image artifacts and coregistration between the optical and MR images (**Fig. 2a**). First, a dental resin wall was made at the circumference of the thinned-skull window, which fit into a single-loop radiofrequency receiver coil for MRI. Second, we immersed an agarose gel dissolved in MRI-invisible D2O to minimize magnetic susceptibility artifacts at the air-skull interface^20^. Third, we introduced enclosed polyethylene (PE) tubes filled with fluorescent rhodamine B (RhB) solution in H2O within the D2O-based agarose gel (**Fig. 2b**). By generating both MR signals and fluorescence, the reference tubes served as landmarks for the coregistration of MR and optical images. For ofMRI studies, we placed the anesthetized mouse onto the MR cradle and adjusted the focal plane of the projection module based on real-time reflectance imaging of the pial vasculature on the cortical surface (**Fig. 2c**).

**Fig. 2.**
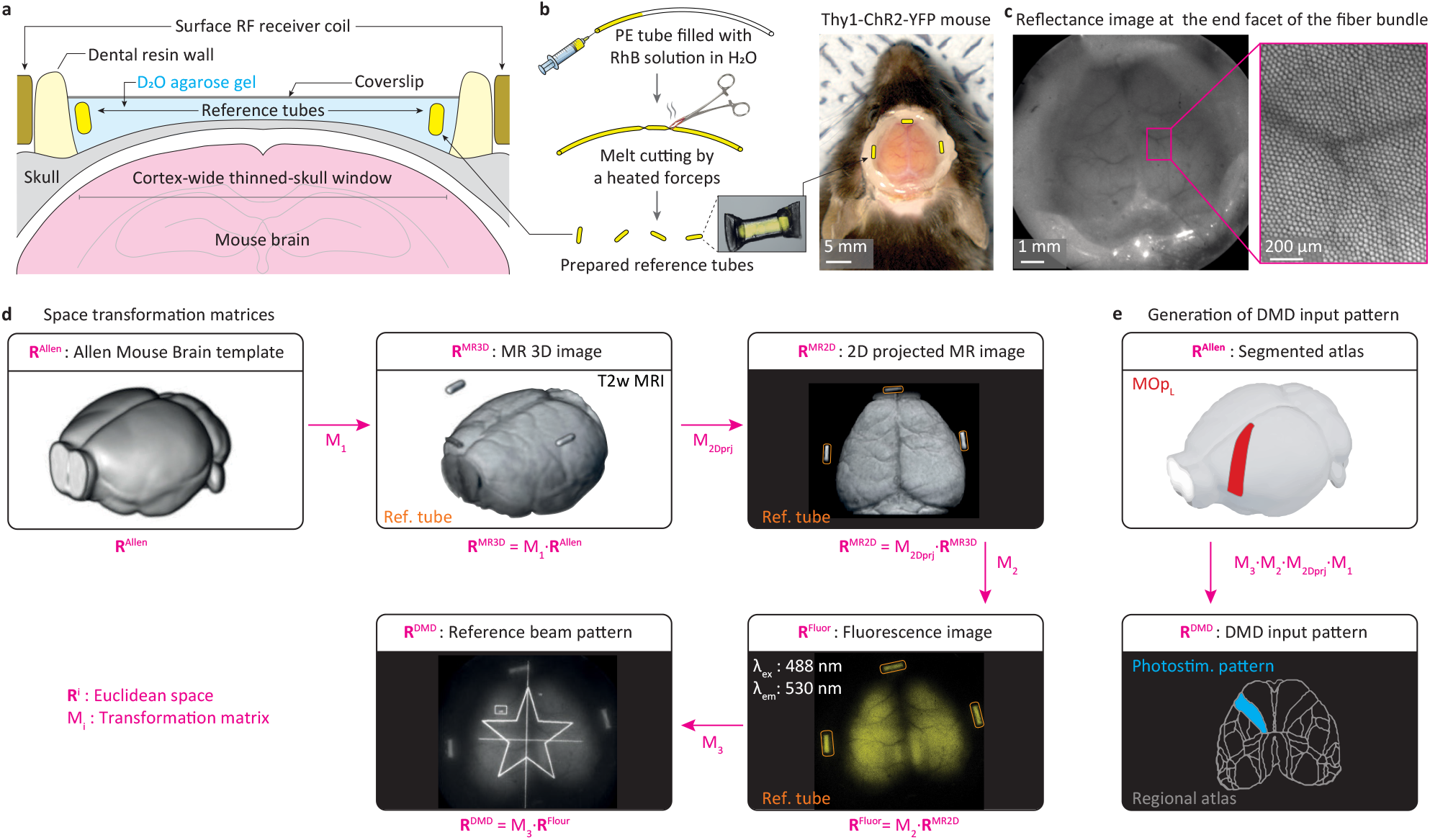
Coregistration of MR and optic spatial coordinates. **a**, Schematic illustration of the cortex-wide thinned-skull window preparation. The cortex-wide thinned-skull window (~1.4 cm in diameter) was prepared on a Thy1-ChR2-YFP mouse. The thinned skull was immersed in a D2O-based agarose gel containing reference tubes for coregistration. **b**, Multiple reference tubes containing rhodamine B (RhB) in H2O solution were prepared by filling the solution inside a polyethylene (PE) tube and melt cutting. The prepared tubes were embedded at the circumference and sealed with a coverslip (right). The cortex-wide blood vessel can be observed by the naked eye after coverslip sealing. A ring shape surface radiofrequency (RF) receiver coil was placed around the dental resin wall (not shown here). Scale bar, 5 mm. **c**, Reflectance image of the agarose gel-covered cortex surface under 530 nm light illumination through a fiber imaging bundle. The pial vascular structure that overlapped with the honeycomb-shaped fiber bundle structure is shown in the enlarged image. Scale bar, 1 mm (left); 500 μm (right). **d**, Derivation of space transform matrices. An anatomical structure-based stimulus map was generated by the following procedures. First, the Allen Mouse Brain template (**R**^Allen^) was transformed into the native MR space (**R**^MR3D^) to obtain M1. Next, the matrix projecting anatomical MR image along the dorsoventral axis (**R**^MR2D^) was defined as M2Dprj. The 2D projected MR image was coregistered to the fluorescence image on the sCMOS camera using the 3 coregistration reference tubes (demarcated by the orange lines), generating M2. Finally, the coordinates of DMD (**R**^DMD^) and the sCMOS camera (**R**^Fluor^) were coregistered by the point-based registration algorithm using a star-shaped polygon pattern on DMD, generating M3. A fluorescence image of the cortical surface was obtained from a transgenic mouse with 488 nm excitation and a 530 nm emission filter set (**R**^Fluor^, YFP channel). **e,** Generation of the DMD input pattern. Using the four transformation matrices (M1, M2, M3, and M2Dprj) in (a), the Allen atlas (**R**^Allen^) was transformed to the DMD space (**R**DMD). This procedure, typically performed within ~30 min, was conducted before every fMRI experiment to generate patterned optogenetic stimuli for individual mice. Scale bar in **b**, 5 mm.

As distinct modalities have their own spatial coordinates, we set a procedure to coregister them into a common coordinate (**Fig. 2d**). The Euclidean spaces involved in our study were the Allen Mouse Brain template (**R**^Allen^), T2-weighted MR images acquired in 3D (**R**^MR3D^) and their 2D projected views (**R**^MR2D^), fluorescence images (**R**^Fluor^), and patterned beam on DMD (**R**^DMD^). Based on the coregistration references, we derived the space transformation matrices, which allowed a one-step generation of the DMD stimulation pattern from the Allen Mouse Brain Atlas (**Fig. 2e**). Using this procedure, we were also able to generate tailored optogenetic stimulation patterns *in situ* based on OIS imaging or a circuit-specific retrograde viral tracer (**Extended Data Fig. 3**).

### Optimization of patterned optogenetic stimuli

Using a previously reported Monte Carlo simulator^21^, we next investigated the volumetric distribution of photons and their photothermal effect in response to patterned optogenetic stimuli. To compensate for the scattering and absorption loss, conventional fiber-optic ofMRI typically involves optical irradiance at the fiber tip greater than the minimally required irradiance for optogenetics (typically ~1 mW/mm^2^)^22^. However, excessive photon density near the tip of the optical fiber potentially causes side effects, such as phototoxicity^23^, photothermal neuromodulation^21,24^, or artifactual fMRI responses^25^. In this simulation, we first compared the two cases, one representing the proposed patterned illumination (beam size = 1 mm, NA = 0.022) and one representing conventional point illumination (beam size = 0.2 mm, NA = 0.22) (**Fig. 3a, b**). Note that point illumination mimics the widely used fiber-optic light delivery approach with a 200 μm core multimode fiber^26–28^. The two different stimulation patterns showed a considerable disparity in terms of penetration depth and tissue temperature. The patterned illumination showed ~2-fold deeper penetration into the cortex than the point illumination, which was due to a larger beam size^29^ (**Fig. 3c**). As expected by the spatial distribution of photons (**Fig. 3a**), point illumination (1 mW, duty cycle: 20%, duration: 10 s) led to the localized accumulation of heat near the cortical surface, while patterned illumination (higher 5 mW, duty cycle: 20%, duration: 10 s) showed a negligible change in tissue temperature (**Fig. 3c**). Our simulation suggests that patterned illumination is advantageous for ofMRI by allowing large numbers of neurons to be targeted with negligible photothermal effects.

**Figure 3.**
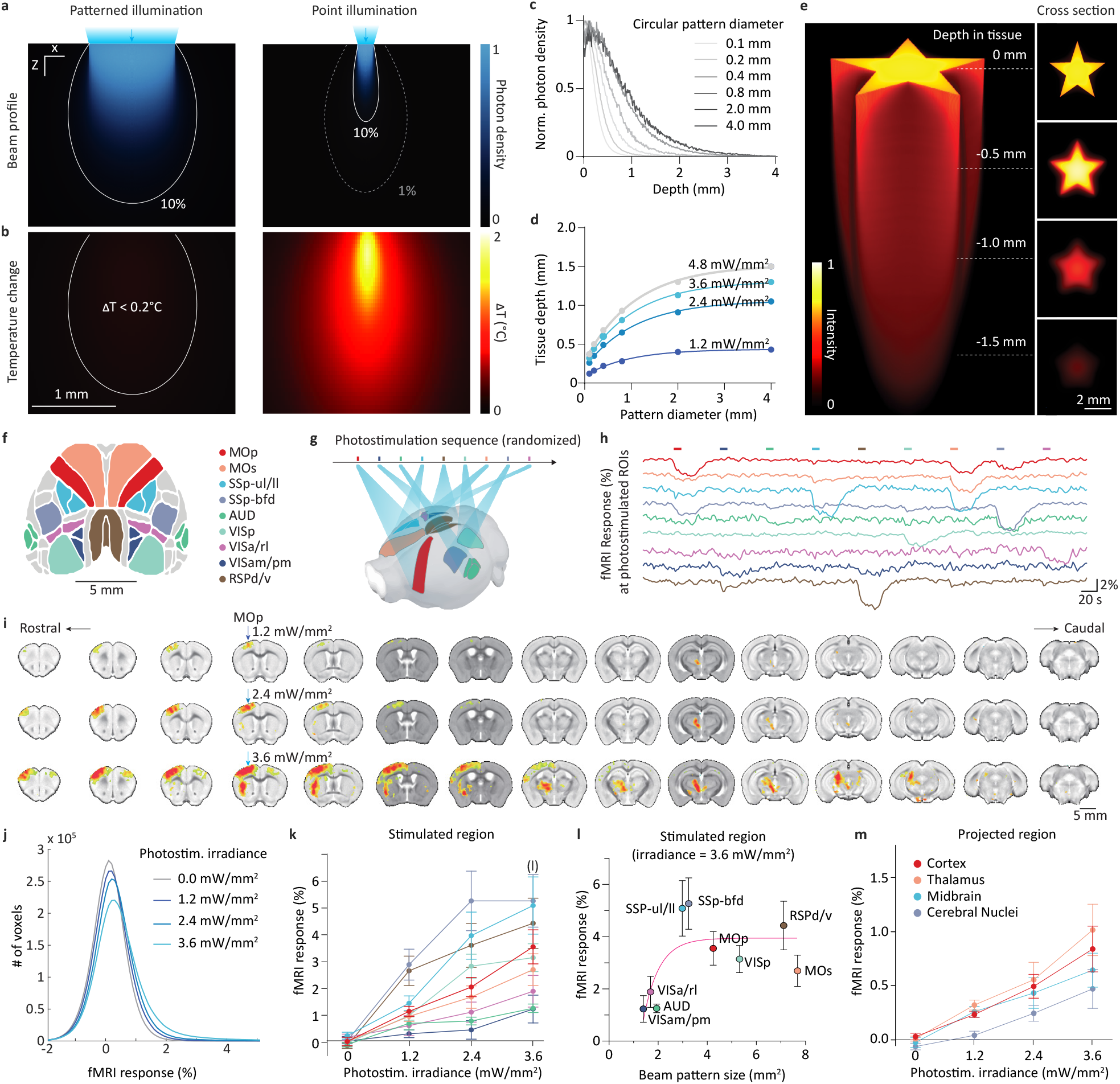
Optimization of patterned irradiance by Monte Carlo simulation and ofMRI in mice. **a-e**, Monte Carlo simulation of beam power distribution in the brain tissue. **a-b**, For simulation, 10-s irradiation with a 20% duty cycle was used for two circular beam conditions, 1-mm diameter with 5 mW (left) and 0.2-mm diameter with 1 mW power (right). Normalized beam power distribution (**a**) and temperature changes (**b**) are shown in the XZ plane section. The 1-mm diameter beam condition with the 5 mW beam heated up the tissue to less than 0.2 °C, but the 0.2-mm diameter beam condition with the 1 mW beam increased the temperature by almost 2 °C. **c**, Depth-dependent light intensity was calculated for different stimulation diameters. **d**, The penetration depth to reach an optogenetically activatable light level of 1 mW/mm^2^ was determined as a function of the beam pattern size and the light intensity. **e**, Spatial deformation of an arbitrarily shaped beam pattern through tissue depth propagation is depicted. Note that the depth axis was elongated only 5 times for visualization purposes. **f-m**, Contrast agent-enhanced, CBV-weighted fMRI studies with patterned optogenetic stimulation in Thy1-ChR2 mice under ketamine/xylazine (K/X) anesthesia. **f**, Cortical ROIs for optogenetic stimulation are defined based on the Allen Mouse Brain Atlas and marked with 9 different colors. **g**, Only unilateral ROIs, except RSPd/v, were used for sequential optogenetic stimulation. MOs, SSp-ul/ll, VISa/rl, and VISam/pm stimuli were applied to the right hemisphere. The order of sequence was randomized per trial. **h**, CBV-weighted fMRI time courses in nine stimulated regions (i.e., the same-colored skeleton regions in **g**) responding to sequential patterned optogenetic stimuli were obtained from one representative mouse. Color bar, 10-s optogenetic stimulus to the same-colored region shown in **g**. Negative changes induced by stimulation over time **(h)** indicate increases in cerebral blood volume. **i**, Coronal fMRI maps showing responses to optogenetic stimulation of the primary motor area (MOp) with three different power levels (top: 1.2 mW/mm^2^; middle: 2.4 mW/mm^2^, bottom: 3.6 mW/mm^2^). Activated volumes increased at the stimulated region (indicated by downward arrows) and projected subcortical regions across the brain. **j,** Averaged fMRI histogram for 4 different optogenetic power levels of 9 stimuli. At a fixed percent change threshold, a larger number of voxels were active in response to higher stimulation power. **k**, Group-averaged fMRI responses in 9 stimulated regions (the same-colored regions in **g**) as a function of stimulus power. **l**, Beam pattern size-dependent fMRI responses to 3.6 mW/mm^2^ irradiance in stimulated regions. A solid line (magenta) indicates nonlinear regression (R^2^ = 0.27). **m**, Irradiance power-dependent group-averaged fMRI responses to 9 optogenetic stimuli in projected regions. All error bars indicate standard errors of the mean.

We next questioned which light irradiance level is optimal for activating cortical neurons, which are distributed to a depth of ~1.3 mm. As expected, higher irradiance led to a deeper effective stimulation depth with a physiological optogenetic irradiance of 1 mW/mm^2^ (**Fig. 3d**). At the typical sizes of the cortical regions of 2–4 mm, we found that an irradiance of 3.6 mW/mm^2^ is optimal for exciting entire cortical layers, without the risk of involving subcortical activation (**Fig. 3d**), and the boundary of the stimulation pattern is largely preserved even in the deep cortical layers (**Fig. 3e**).

To verify our simulation results with *in vivo* studies, we performed ofMRI in Thy1-ChR2 mice under light anesthesia with three different irradiance levels on nine different atlas-based cortical regions: unilateral motor (primary, MOp; secondary, MOs) areas, primary somatosensory (barrel field, SSp-bfd; upper and lower limb, SSp-ul/ll) areas, visual (primary, VISp; anterior and rostrolateral, VISa/rl) areas, auditory (AUD) areas, medial (antero- and posteromedial visual, VISam/pm) areas and bilateral retrosplenial areas (dorsal and ventral, RSPd/v)^30^ (**Fig. 3f-m**). Instead of conventional blood oxygenation level-dependent (BOLD) fMRI, cerebral blood volume (CBV)-weighted fMRI with 200 μm isotropic resolution was adopted with an intravenous bolus injection of long-half-life blood-pool iron oxide nanoparticles to enhance functional sensitivity and specificity^22^. Each fMRI run consisted of nine blocks with a 10 s optogenetic stimulus (20 Hz, 20% duty cycle) and 50 s recovery periods in a semirandomized sequence of stimulation regions (**Fig. 3g**, **Supplementary Video 1**). In the raw time traces on targeted regions of interest (ROIs) (i.e., same-colored time courses as stimulation), anatomy-based optogenetic stimuli reliably evoked increases in CBV (corresponding to decreases in fMRI signals) in the pertinent ROIs, suggesting the effectiveness of patterned photostimulation (**Fig. 3h**). The magnitude and extent of the fMRI responses increased with increasing irradiance power (**Fig. 3i, j**), but the magnitudes in the stimulated sites were heterogeneous (**Fig. 3k**), presumably due to the influence of the beam size (**Fig. 3l**) and different neurovascular coupling. Overall, the irradiance of 3.6 mW/mm^2^ provided sufficient functional sensitivity for all nine targeted and projected regions (**Fig. 3k, m**) and thus was chosen for subsequent fMRI studies.

### Brain-wide comparison of structural connectivity (SC) and EC

Having established that our patterned ofMRI robustly measures the causal influence from the targeted cortical regions to other brain areas, we next investigated the relationship between large-scale functional networks and structural connections, which is fundamental for linking structure to function^31,32^. We acquired EC data with cortex-wide patterned optogenetic stimulation of the nine cortical regions (MOp, MOs, SSp-bfd, SSp-ul/ll, VISp, VISa/rl, AUD, VISam/pm, and RSPd/v) in anesthetized Thy1-ChR2 mice (n = 8 mice with ketamine/xylazine), resulting in robust activation in the photostimulated and networked regions, mainly on the ipsilateral side (**Extended Data Fig. 4**). For SC, we used the Allen Mouse Connectivity Atlas (https://connectivity.brain-map.org) constructed with anterograde viral tracers focally injected into the same nine cortical regions, visualizing brainwide monosynaptic axonal projections originating from the injected sites (**Fig. 4a, b**).

**Fig. 4.**
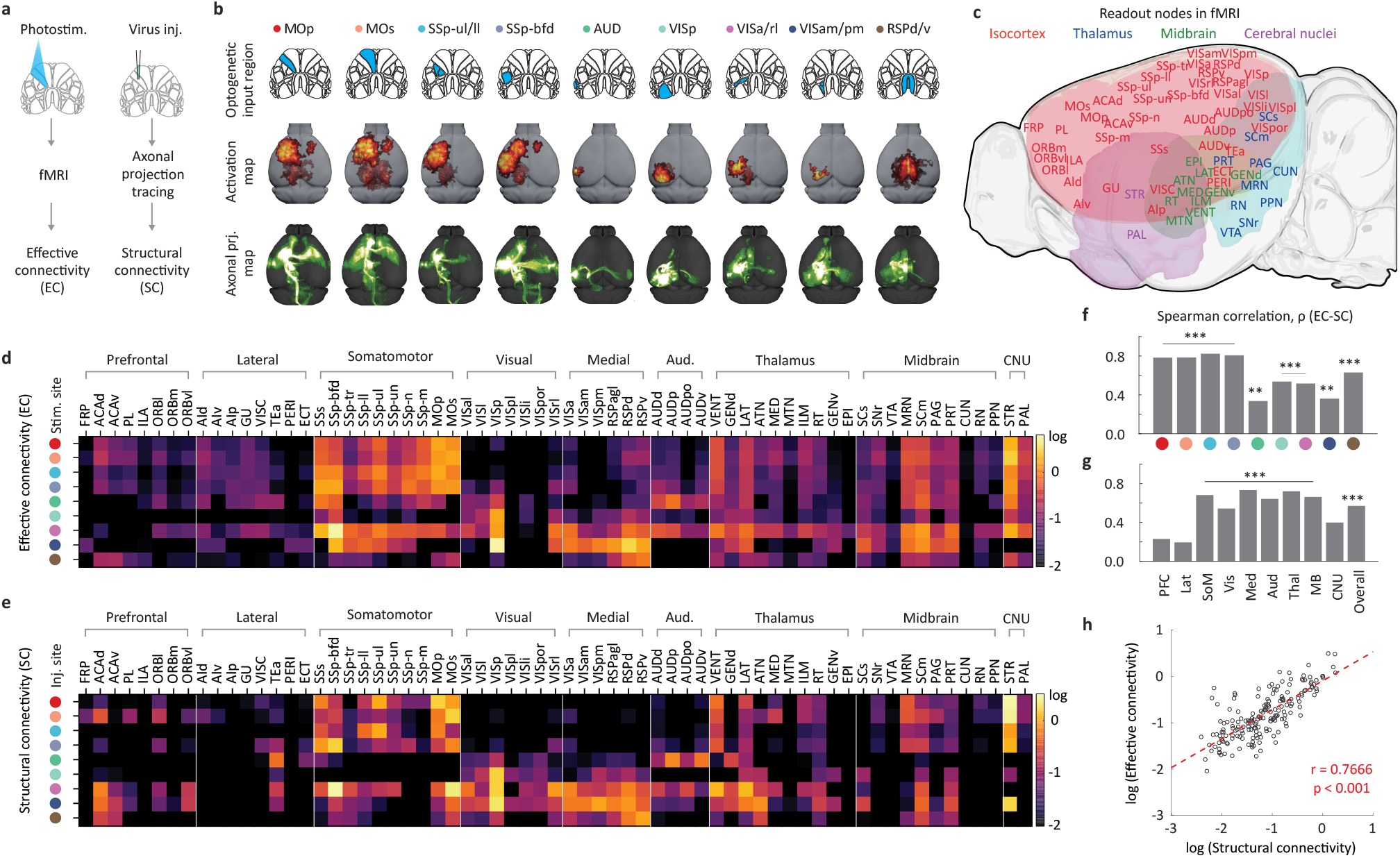
EC measured by brain-wide fMRI with patterned optogenetic stimuli vs. SC. **a**, Schematic of the procedures for acquiring EC and SC. **b**, fMRI and axonal connectivity maps. The color code indicating each source (stimulus or injection site) is used throughout this figure. (Top) Optogenetic illumination patterns used for EC mapping in fMRI. (Middle) The dorsal view of group-averaged fMRI activation maps (familywise error rate (FWER)-corrected p<0.05, n = 8 mice). (Bottom) Axonal projection patterns imported from the Allen Mouse Brain Connectivity Atlas (http://connectivity.brain-map.org; Experiment # 127084296, 112952510, 112229814, 112951804, 112881858, 307593747, 657334568, 100141599, and 100140949). Note that bilateral RSPd/v stimulation was performed in fMRI experiments, while the corresponding axonal projection was identified by unilateral injection. The left side of the brain was the hemisphere ipsilateral to the stimulation site. **c**, Target ROI definition for quantitative analyses. Sixty-five ipsilateral ROIs from the isocortex, thalamus, midbrain, and cortical nuclei were defined. **d, e**, Comparison of EC (**d**) and SC (**e**) from 9 sources (rows) to 65 targets (columns) in nine network modules. Responses with negative EC values, mostly spurious (except for RSPd/v-VISpl, see Fig. 5b for statistical analysis), were considered not to be active. The SC datasets from the Allen connectivity atlas were modified to conform with the fMRI dataset (see Methods). The connection strengths are on a log scale. **f, g,** Spearman correlation coefficient between EC and SC from each source to all ROIs (**f**) and for all 9 sources to all ROIs within the same network module (**g**). **, p<0.01; ***, p<0.001. **h**, Comparison of EC and SC strengths. Both connection strengths are highly correlated, yielding a Pearson correlation coefficient of 0.7666.

For quantitative comparison, we defined 65 atlas-based ipsilateral ROIs in the isocortex, thalamus, midbrain, and cerebral nuclei (**Fig. 4c**; **Supplementary Table 1**), which were further classified into 9 network modules based on the network modularity of the SC^33^. Then, we calculated EC strength as the sum of voxelwise fMRI responses in each ROI of all animals divided by the sum of responses in the stimulated ROIs of all animals and we defined SC strength as the fluorescence pixel counts of each ROI normalized to those of the source ROI. The EC and SC data from the 9 cortical source regions (i.e., photostimulation for EC and virus injection for SC) to the 65 target ROIs over the brain are presented as heatmaps (**Fig. 4d, e**). Overall, the measured EC data greatly resembled the SC dataset. All the EC data for each cortical source region (i.e., each row in the EC heatmap) showed significant positive correlations with the corresponding SC data (Spearman coefficient, ρ = 0.57; p < 0.001; **Fig. 4f**). Notably, AUD and VISam/pm showed a lower correlation than other cortical source regions (ρ ~ 0.3), conceivably due to the lower fMRI responses in the photostimulated regions (see **Fig. 3l**). When target network modularity was compared between EC and SC data, 6 modules (among 9), including somatosensory, visual, medial, auditory, thalamic, and midbrain modules, showed significant positive correlations (p < 0.001; **Fig. 4g**).

The similarities and differences between SC and EC were further examined with binary matrices (**Extended Data Fig. 5a-c**), where the existence of EC was determined by a statistical test across subjects (one-sample t-test; false discovery rate-corrected p<0.05). The binarized EC and SC matrices showed a high congruency of 66% (n = 188 out of 9 × 65 = 585 connections for ‘both’ and n = 197 for ‘neither’; **Extended Data Fig. 5a**), although the congruency was dependent on a threshold level of SC strengths and the stimulated region (**Extended Data Fig. 5b-c**). The congruent connection strengths for ‘both’ (n = 188 connections) showed a highly positive linear correlation (Pearson’s r = 0.767) (**Fig. 4i**). A similar relationship between EC and SC was also observed under isoflurane (ISO) anesthesia (n = 9 mice with ISO, **Extended Data Fig. 5d-f, Extended Data Figs. 6-7**). Overall, the results of our EC vs. SC investigation suggested that the measured EC largely reflects the monosynaptic axonal projections.

### Modulation of brain-wide EC by anesthetics

Anesthesia can affect neural processing at the local level by changing the balance of excitatory and inhibitory activities and can consequently act in a circuit-specific manner^34,35^. Thus, we sought to identify differences in brain-wide EC between two widely used anesthetics, ketamine/xylazine (K/X) and ISO (**Fig. 5a**). fMRI responses to nine optogenetic stimuli were obtained under either N-methyl-D-aspartate (NMDA) antagonist ketamine and α2-adrenergic agonist xylazine (K/X, n = 8 mice) or potent gamma aminobutyric acid A (GABA_A_) agonist isoflurane (ISO, n = 9 mice) (**Fig. 5b, c** for significant EC regions by one-sample t-test with false discovery rate (FDR)-corrected p < 0.05). In general, ISO reduced the magnitudes of functional responses relative to K/X, which may be due to different neurovascular coupling properties under different anesthetics^15^. Thus, we compensated for this neurovascular coupling variability **(Fig. 5d)** by normalizing each subject’s responses to the mean response of nine photostimulated source sites. Then, the normalized ISO responses were subtracted from the normalized K/X responses, yielding a difference matrix (**Fig. 5e**; Mann–Whitney U test; FDR-corrected p<0.1 were marked as asterisks). Overall, ISO reduced responses in circuits from cortical sources to the somatomotor, thalamus, and midbrain areas but enhanced responses in visual areas.

**Fig. 5.**
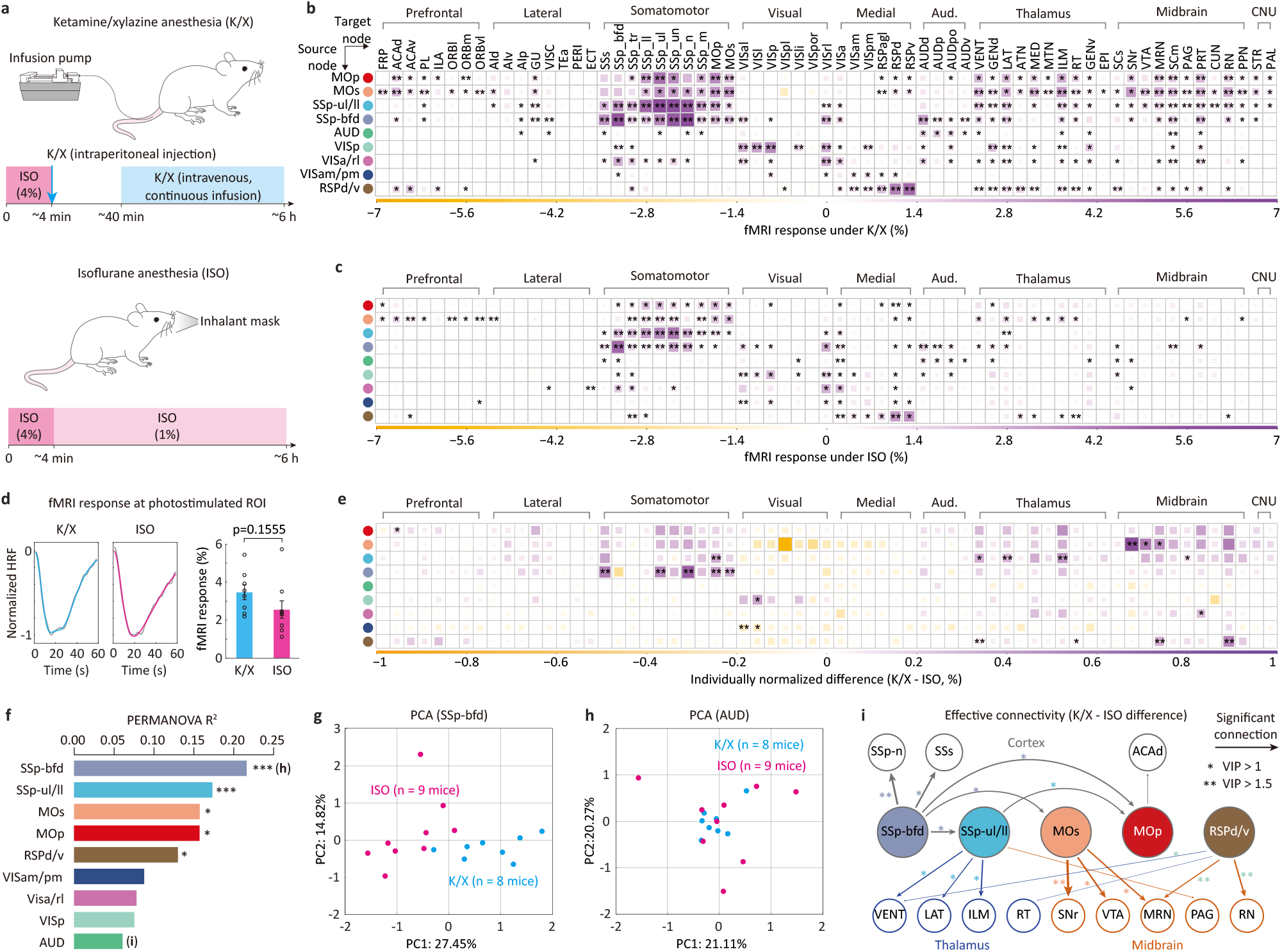
Brain state-dependent EC changes. **a**, Anesthetic protocol for K/X continuous infusion and ISO inhalation. CBV-weighted fMRI with nine anatomically defined optogenetic stimuli was performed in 8 K/X- anesthetized and 9 ISO-anesthetized Thy1-ChR2 mice. **b-c**, Group-averaged response matrix for K/X (**b**) and ISO anesthesia (**c**). A one-sample t-test with FDR correction was performed for statistical analysis (*, p<0.05; **, p<0.01). **d**, Normalized hemodynamic response function (HRF) for the K/X and ISO groups (left panels) and animalwise responses averaged in the 9 stimulated ROIs (right panel). No significant difference was observed between K/X and ISO conditions (independent t-test). fMRI responses were normalized by the mean responses of stimulation sites in each subject. **e**, A difference matrix of the normalized responses between the K/X and ISO groups (*, p<0.1; ** p<0.05, Mann–Whitney U test with FDR correction). **f**, PERMANOVA showed the explained variance (R^2^) of each stimulus (*, p<0.05; **, p<0.01; ***, p<0.001). **g, h**, PCA of SSp-bfd and AUD stimulation-induced response matrix (each contains 17 mice x 65 target ROIs). The individual data points projected onto principal components (PCs) 1 and 2 are well differentiated as ISO and K/X groups for SSp-bfd stimulation but not for AUD stimulation. **i**, Significantly reduced EC under ISO conditions compared to that under K/X conditions, confirmed by a two-staged validation procedure. The connections with variable importance in projection (VIP) values greater than 1 were considered important in the partial least square model (asterisks).

To identify which stimulus source produces discriminable fMRI responses under different anesthesia conditions, we performed permutational multivariate analysis of variance (PERMANOVA; **Fig. 5f**)^36^ for each stimulus-induced response matrix (17 animals x 65 ROIs). Stimulation of SSp-bfd, SSp-ul/ll, MOs, MOp, and RSPd/v provided significantly differentiated responses under K/X vs. ISO conditions. The PERMANOVA results were further examined by principal component analysis (PCA; **Fig. 5g, h**); two anesthetic groups were clearly separated upon the first and second principal components of the SSp-bfd EC matrix (the highest explained variance R^2^ of 0.22) but this was not the case with AUD EC (the lowest R^2^ of 0.06). Responses to the five cortical stimulation sites were significantly weakened under ISO in eighteen target regions (**Fig. 5i**). The relative significance of each connection in discriminating anesthetic conditions was assessed by the partial least square-discriminant analysis (PLS-DA) approach, yielding variable importance in projection (VIP) values (asterisks in **Fig. 5i**; **Extended Data Fig. 8** for details). SSp-bfd to SSp-n (nose region in the primary somatosensory area), MOs to SNr (the reticular part of the substantia nigra), and RSPd/v to the midbrain reticular nucleus (MRN) and red nucleus (RN) connections (VIP > 1.5) were the most significantly altered by ISO. This demonstrates that whole-brain fMRI with patterned optogenetics can measure changes in EC induced by brain status.

## DISCUSSION

In this study, we reported a novel hybrid system that incorporates cortex-wide patterned photostimulation in ofMRI. By providing highly flexible delivery of optogenetic stimuli in space and time, we obtained an unprecedented comprehensive map of EC originating from cortical regions, which showed high similarity with the connections shown in the Allen Mouse Connectivity Atlas. We further applied our system to dissect the anesthesia-dependent modulation of specific neural circuits, implicating its diverse applications in studying altered functional brain states in psychiatric and neurodegenerative diseases^37–40^.

By taking advantage of both optics and MRI, as well as genetics, our patterned ofMRI has versatile applications. For individual subject-specific studies, photostimulation patterns can be generated from *in situ* optical or MRI. To demonstrate this capability, cortical activation sites responding to forepaw stimulation were simultaneously measured by intrinsic optical imaging and fMRI, and optogenetic stimulation of the active cortical region induced brain-wide functional activity largely resembling forepaw stimulation (**Extended Data Fig. 3a, Supplementary Video 2**). Similarly, circuit-specific studies can be performed by generating photostimulation patterns based on fluorescent neurons labeled by anterograde or retrograde tracers. As a proof-of-principle, we injected a retrograde viral vector encoding ChR2 with a yellow fluorescent protein at the inferior colliculus (IC) and then photostimulated the IC-projecting neurons in the AUD cortex defined by fluorescent images (**Extended Data Fig. 3b**). Alternatively, cell-type-specific EC can also be obtained by activating genetically specified neuronal subtypes, such as inhibitory neurons, using VGAT-ChR2 mice (**Extended Data Fig. 9**). In addition to functional mapping in the spatial domain, interactions between multiple cortical areas may be investigated in the time domain at a millisecond scale; as an example, we observed brain-wide functional modulation driven by synchronous and delayed bilateral photostimulation on the right and left S1 (**Extended Data Fig. 10**)^41^.

There are technical limitations of DMD-based patterned optogenetics. First, since the tissue penetration depth is limited to less than 1.5 mm, it is only feasible for photostimulation of cortical regions. Although introducing longer-wavelength or ultrasensitive optogenetic actuators may allow greater effective depths, conventional fiber-optic light delivery is recommended to target subcortical areas^42,43^. Second, in contrast to fiber-optic light delivery, remote targeting of an underlying structure is not feasible. For example, our current system does not support targeting layer 4-5 cortical neurons without affecting the superficial neurons. This issue may be mitigated genetically by using layer-specific targeted expression^44^. Third, as shown in our optical simulation, the spatial profile of the effective photostimulated area is dependent on the size of the targeted regions. This effect is noticeable if the photostimulation area is small (< 1 mm^2^), leading to significantly lower fMRI responses in the photostimulated and networked regions (e.g., AUD and VISam/pm). This issue can be solved by adjusting the optical irradiance depending on the stimulation area.

With our newly developed methodology, we investigated the relationship between FC and SC, which has been a major focus of systems neuroscience research^45,46^. Conventional FC measured by fMRI relies on covaried hemodynamic modulations at a resting state (RS). However, the underlying sources of RS FC are unclear and do not faithfully reflect neuronal RS circuits in mice^47^. Thus, we adopted EC measurements by directly modulating neuronal activities in source regions and reading out corresponding responses in the entire brain by fMRI. Interestingly, our EC data under anesthesia are largely restricted to monosynaptic axonal connections, which is in agreement with optical intrinsic imaging-based EC mapping in the cortex^13^. However, 30-40% incongruency was observed with the SC dataset. The incongruency is found throughout the brain, where SC-only components (25% and 38% for K/X and ISO, respectively) are more predominant than EC-only components (9% and 3%). The inconsistency between SC and EC is likely due to several reasons: highly sensitive SC is monosynaptic and has limited injection sites, whereas hemodynamic-based EC can be polysynaptic and has relatively low sensitivity. Moreover, the use of anesthesia would lead EC suppression in a nonspecific and circuit-specific manner^48^. To further investigate the anesthetic effect on EC, a protocol for awake mice can be implemented for patterned optogenetic fMRI^49^.

To conclude, our method expands the flexibility and productivity of ofMRI. We demonstrated the high-throughput capability for obtaining multisite EC for the construction of an “effective functional connectome”. Note that the functionality of our approach can be tremendously expanded when integrated with numerous optogenetic lines, in keeping with attempts to reveal cell-type-specific axonal projection patterns^33^. Moreover, since our optic module can be easily assembled with commercially available parts and integrated with existing preclinical MR systems, we envisage that our method can be readily adopted in a variety of fMRI studies that require i) sequential cell-type-specific stimulation on multiple sites for mapping functional cortico-cortical or subcortical networks, ii) temporally controlled stimulation on multiple sites for investigating interregional functional coupling balances, iii) an *in situ* subject-specific stimulation design defined by anatomical or functional imaging, and iv) simultaneous functional mapping of cortical areas through optical imaging and cortical/subcortical regions by fMRI. Our method opens new opportunities for investigating dynamic brain statedependent (e.g., plasticity, disease progression, and therapeutic interventions) circuit dissection or biomarker screening in whole-brain functional networks.

## Supporting information

Supp_Video1_Experimental scheme

Supp_Video2_OIS

## Acknowledgments

This research was supported by the Institute of Basic Science (IBS-R015-D1) and the Basic Science Research Program through the National Research Foundation of Korea (NRF), funded by the Ministry of Education (2019R1C1C1011180, 2019M3A9E2061789, 2019M3E5D2A01058329, 2020M3C1B8016137, and 2020R1A5A1018081). We thank Haiyan Jiang for performing viral injections.

## Author contributions

S.K., M.C., and S.-G.K. initiated and supervised the study. S.K. designed and implemented the optical system. S.K., T.T.V., and H.S.M. performed the experiments and data analysis. H.S.M performed MR image processing and data analysis. G.H.I. controlled and optimized the physiological condition of the animals during the experiments. S.K., H.S.M., M.C., and S.-G.K. wrote the manuscript.

## Corresponding author

Correspondence to Seong-Gi Kim and Myunghwan Choi

---

**Extended Data Fig. 1.**
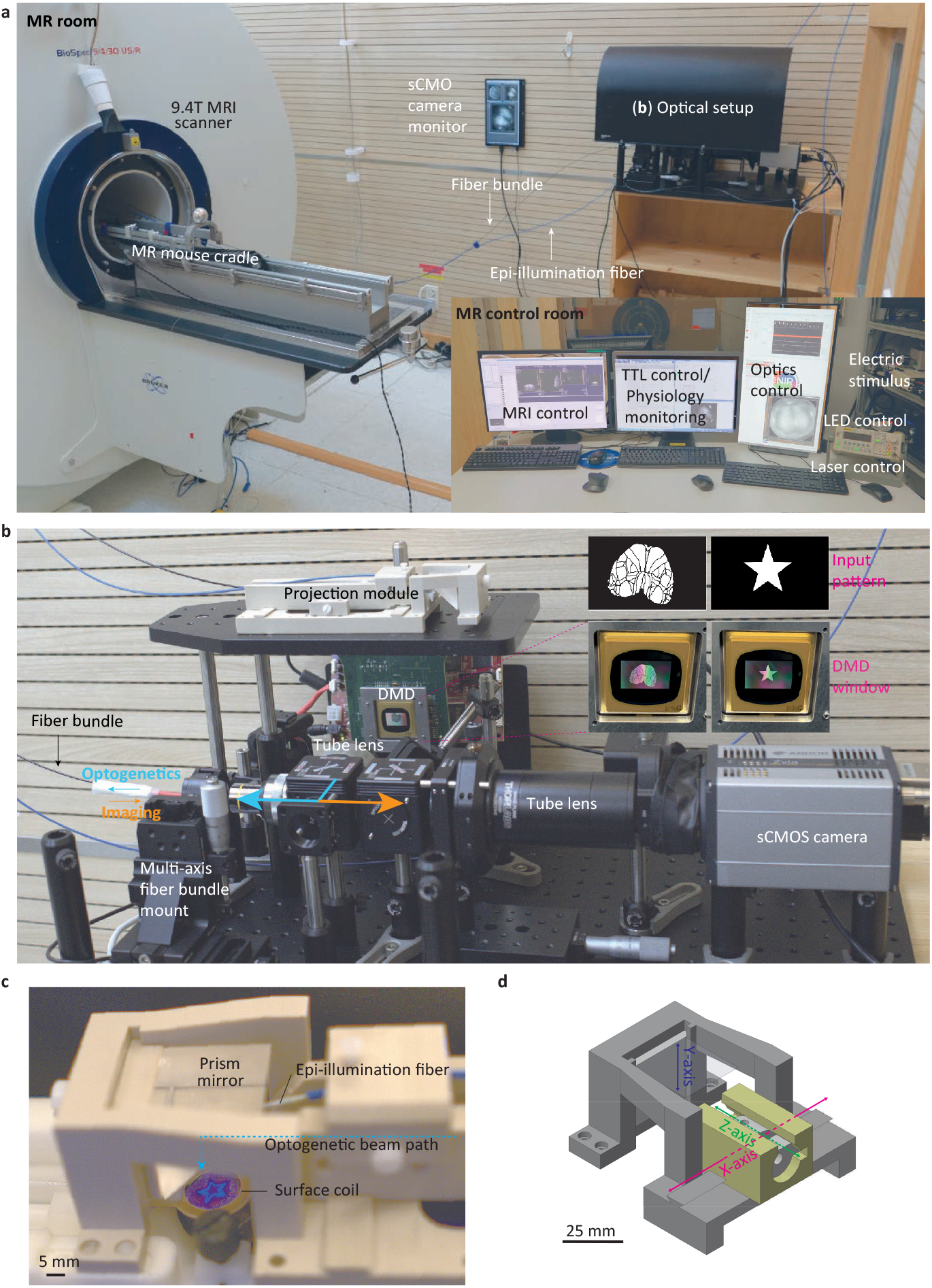
Photographs of the overall patterned ofMRI system. **a**, A picture of the ofMRI setup in the MR room. The external optical setup (marked (b) in the right upper corner) is affixed at the corner of the 9.4T MR room. An LCD monitor on the wall displays reflectance or fluorescence images of the cortical surface within the MR bore. In the MR control room located next to the MR room (inset picture), imaging and physiological data were recorded, and all stimuli were controlled. **b**, Optical setup inside the MR room. The setup combines the imaging path (from the fiber imaging bundle to the sCMOS camera; orange arrow) and the illumination path (from the DMD to the fiber bundle; cyan arrow) via a dichroic filter cube. The proximal tip of the fiber bundle is focused on the objective lens with XYZ and rotational stages. The DMD window and the camera sensor planes are optically conjugated. The projection module, which is attached to the mouse MR cradle, is usually kept on the optical board. Computer-generated binary input pattern images and their appearance on the DMD window are shown as insets. **c**, The projection module is mounted over the head of the mouse. The epiillumination fiber was attached on top of the projection module. The surface coil was placed on the mouse around the dental resin wall. A star-shaped beam pattern was projected onto the mouse brain for demonstration purposes. **c**, Kinematic optical mount in the projection module. By sliding parts of the module, the position of the beam pattern and imaging field of view can be adjusted.

**Extended Data Fig. 2.**
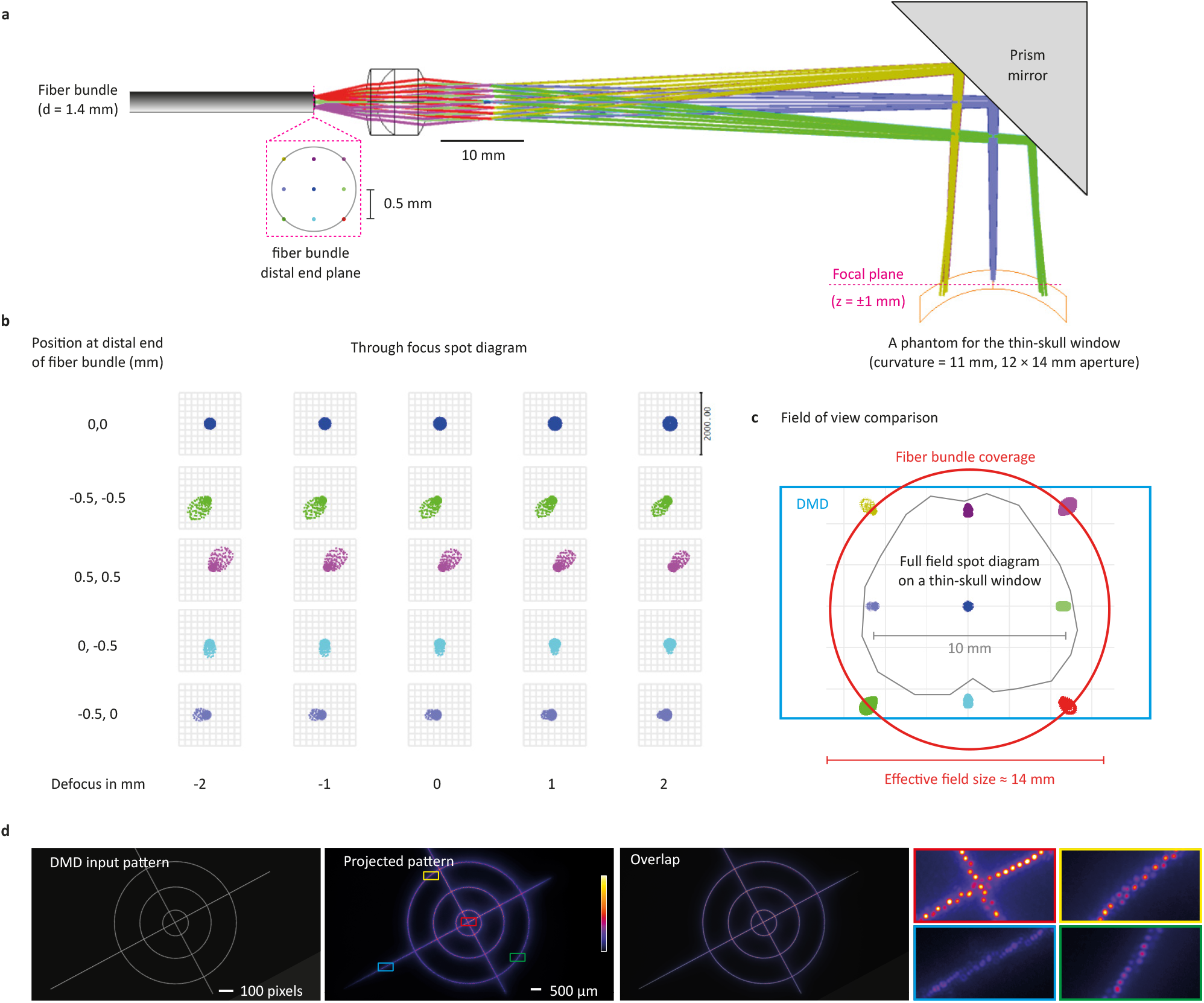
Optical design of the projection module. **a.** Zemax simulation of the optogenetic beam pattern projected on the cortex area. Beam rays are emitted from a 3×3 grid array from the distal end of the surface of the bundle fiber with a 0.5 mm distance and 0.22 NA. Beam rays are reflected and projected on a curved mouse cortical window phantom. **b**. Representative beam spot diagram through focus showing focal plane change-insensitive beam spot distortion within 1~2 mm of the estimated focal depth plane changes in the thinned-skull window. **c**. Field of view (FOV) coverage of the fiber bundle and DMD active window for a thinned-skull window is relatively illustrated. **d**. A projected optogenetic pattern was acquired by placing a CMOS camera (MU895, Thorlabs) at the focal plane. The projected pattern reflected the shape of the DMD input pattern well. The individual cores of a fiber bundle were clearly distinguished from the expanded images.

**Extended Data Fig. 3.**
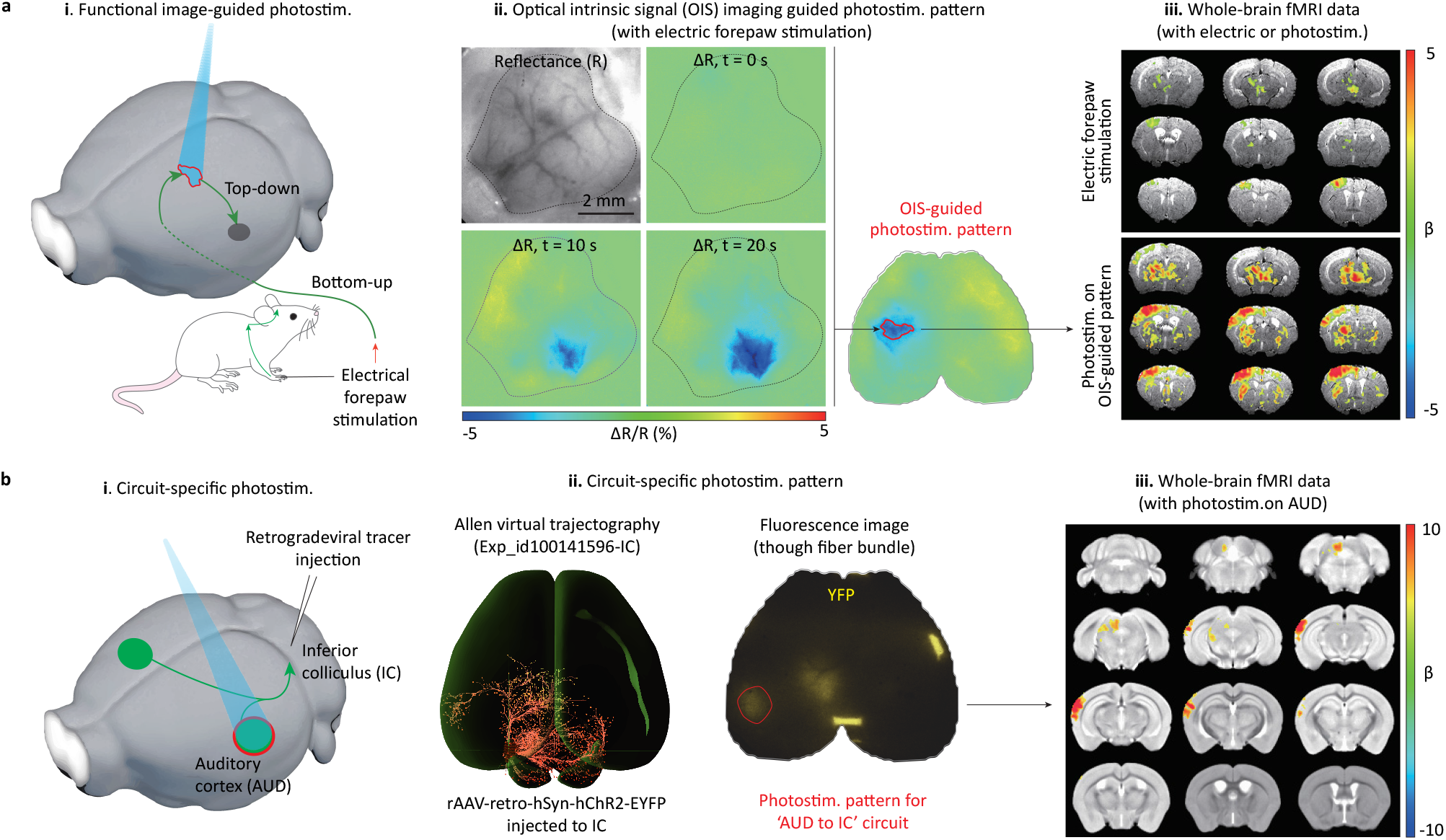
Demonstration of *in situ* patterned optogenetic mouse fMRI. **a**, An optogenetic stimulation pattern was extracted from hemodynamic response images via OIS under forepaw electric stimulation (i). During the 20-s forepaw stimulation, a decrease in OIS signals was observed, which is indicative of an increase in blood volume (ii). ΔR/R, relative reflectance change in total hemoglobin-weighted visible light. The OIS-extracted pattern (at time = 10 s after the onset of stimulation) was converted to the optogenetic stimulation pattern and projected onto the responsive region of the mouse cortex. fMRI activation maps obtained by regional optogenetic stimulation and electric stimulation were highly overlapped (iii). **b**, Circuit-specific activation was demonstrated by illuminating an optogenetic pattern on the region identified by fluorescence imaging (i). Circuit-specific cells projecting from the AUD cortical region to the IC subcortical region were identified by yellow fluorescent images (ii) after viral injection of rAAV-retro-hSyn-hChR2-EYFP into the IC. The circuit-specific activity (AUD -> IC) was measured via fMRI readout by photostimulation of circuit-specific cells in the AUD region (iii).

**Extended Data Fig. 4.**
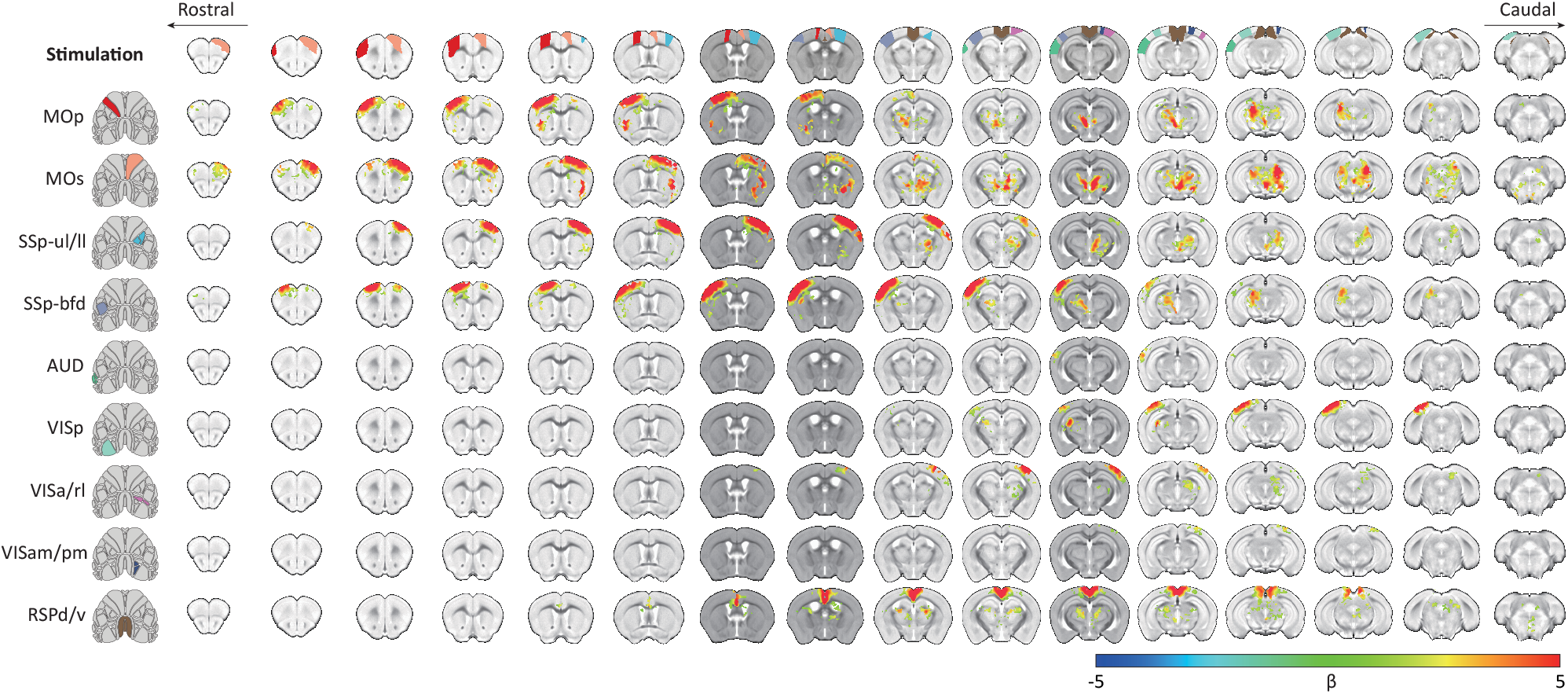
Group-averaged fMRI activation maps of the K/X group (n = 8 mice) by photostimulation of 9 atlas-based regions. Optogenetic stimulus patterns are shown in the left panel, and the locations of stimulation ROIs are depicted at the top. Voxels with FWER-corrected p<0.05 were considered significant.

**Extended Data Fig. 5.**
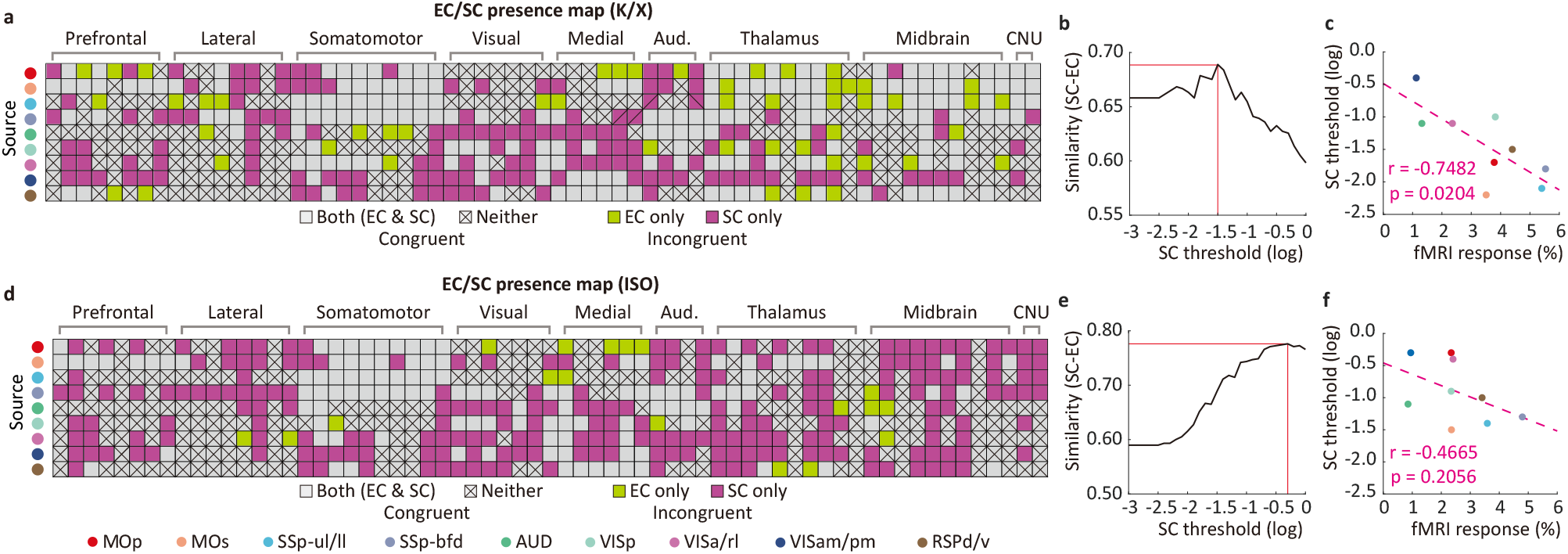
Similarity between EC and SC as a function of the SC threshold. The same procedure was conducted on the K/X (**a-c**) and ISO datasets (**d-f**). **a**, **d**, Comparison of EC and SC binary matrices. The presence of EC was determined by a statistical test across animals, and SC was thresholded at a projection volume of 0.008 mm^3^ corresponding to the volume of one fMRI voxel. **b**, **e**, Based on the assumption that SC-only connections are due to low fMRI sensitivity, the similarity between SC and EC (1 – Hamming distance) was computed as a function of the SC threshold, which is maximized at −1.5 under K/X and −0.3 under ISO. This indicates that the general detectability is reduced under ISO. **c, f**, The optimal SC threshold was also computed with each stimulus (each row in **a, d**). The optimal SC threshold is negatively correlated with fMRI responses at the stimulation site, suggesting that a high fMRI response to optostimulation is needed for mapping EC.

**Extended Data Fig. 6.**
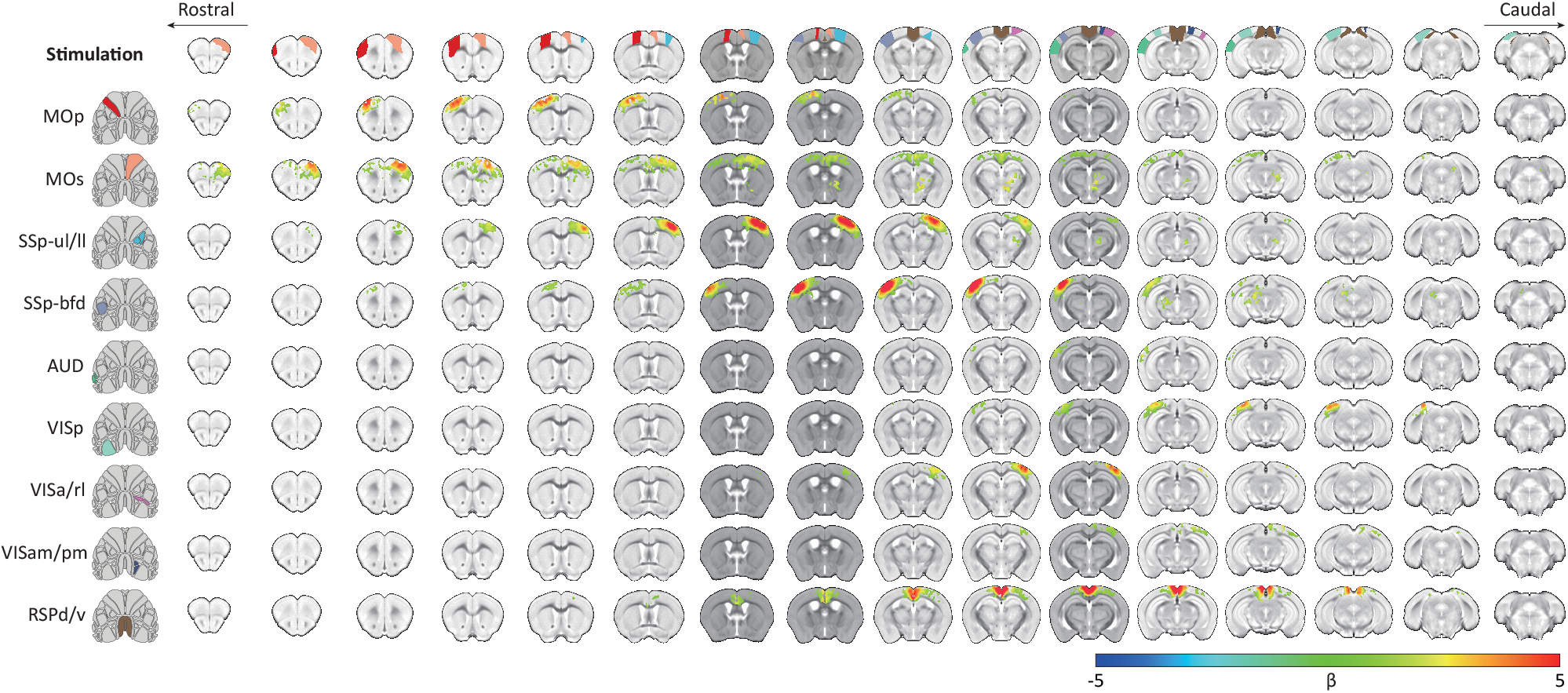
Group-averaged fMRI activation maps of the ISO group (n = 9 mice) by photostimulation of 9 atlas-based regions. Optogenetic stimulus patterns are shown in the left panel, and the locations of stimulation ROIs are depicted at the top. Voxels with FWER-corrected p<0.05 were considered significant.

**Extended Data Fig. 7.**
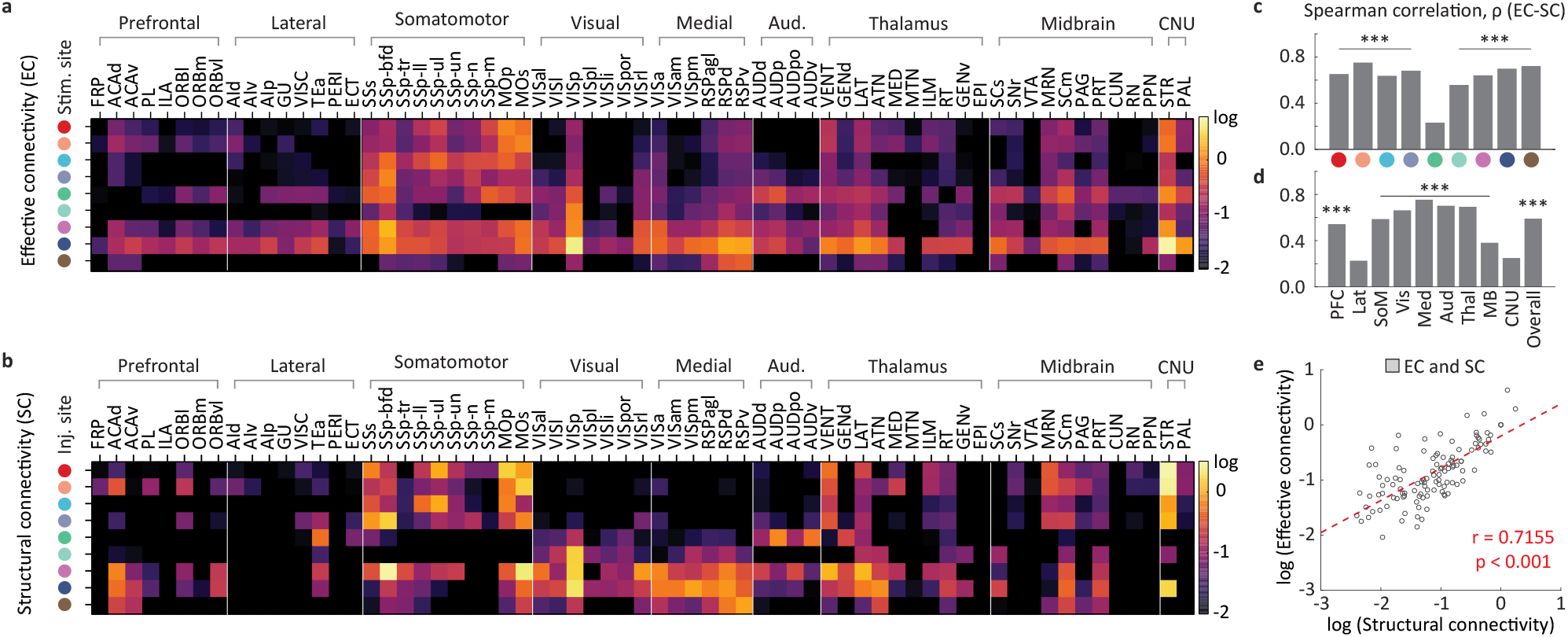
Correspondence between SC and EC under ISO. **a, b**, Comparison of EC (**a**) under ISO and SC (**b**). Responses with negative EC values, mostly spurious (except for RSPd/v-RN), were considered not significant. The SC datasets in b are the same as those in Fig. 4e. Each row represents the stimulation source site, and each column represents the readout target ROI. The connection strengths are on a log scale. **c, d,** Spearman correlation coefficient between EC and SC for each source for a total of 65 ROIs (**c**) and for all 9 sources for all ROIs within the same network module (**d**). ***, p<0.001; *, p<0.05. **e**, Comparison of EC and SC strengths for congruent EC and SC connections. Both connection strengths are highly correlated, yielding a Pearson correlation coefficient of 0.7155.

**Extended Data Fig. 8.**
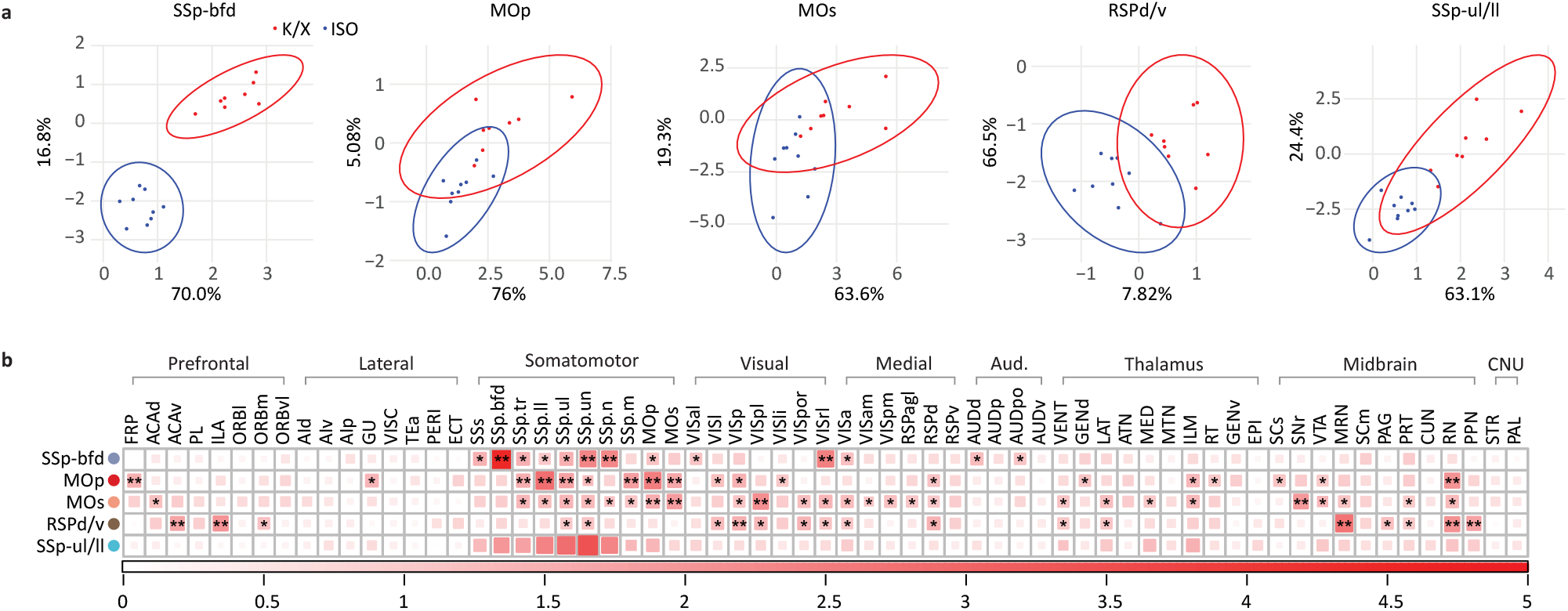
Discriminative networks to differentiate K/X and ISO anesthetic actions. EC data from five cortical source regions, proven to offer a significant difference between K/X and ISO connectivity (**Fig. 5g**), were further examined to assess the contribution of each connection in discriminating anesthetic conditions. **a,** PLS-DA was performed with individual response matrices in separate groups (n = 8 mice for K/X; n = 9 mice for ISO) for each source region. Score maps for individuals are presented upon the first two PLS component axes, where percentage values represent the explained variances by each PLS component. For each source region, R2Y/Q2Y were calculated as follows: 0.949/0.934 for SSp-bfd; 0.683/0.466 for MOp; 0.575/0.501 for MOs; 0.741/0.263 for RSPd/v; and 0.742/0.699 for SSp-ul/ll. For Q2Y, fivefold cross-validation was performed. **b,** The degree of meaningfulness of each connection was evaluated by a VIP score heatmap for PLS-DA (*, VIP>1; **, VIP>1.5).

**Extended Data Fig. 9.**
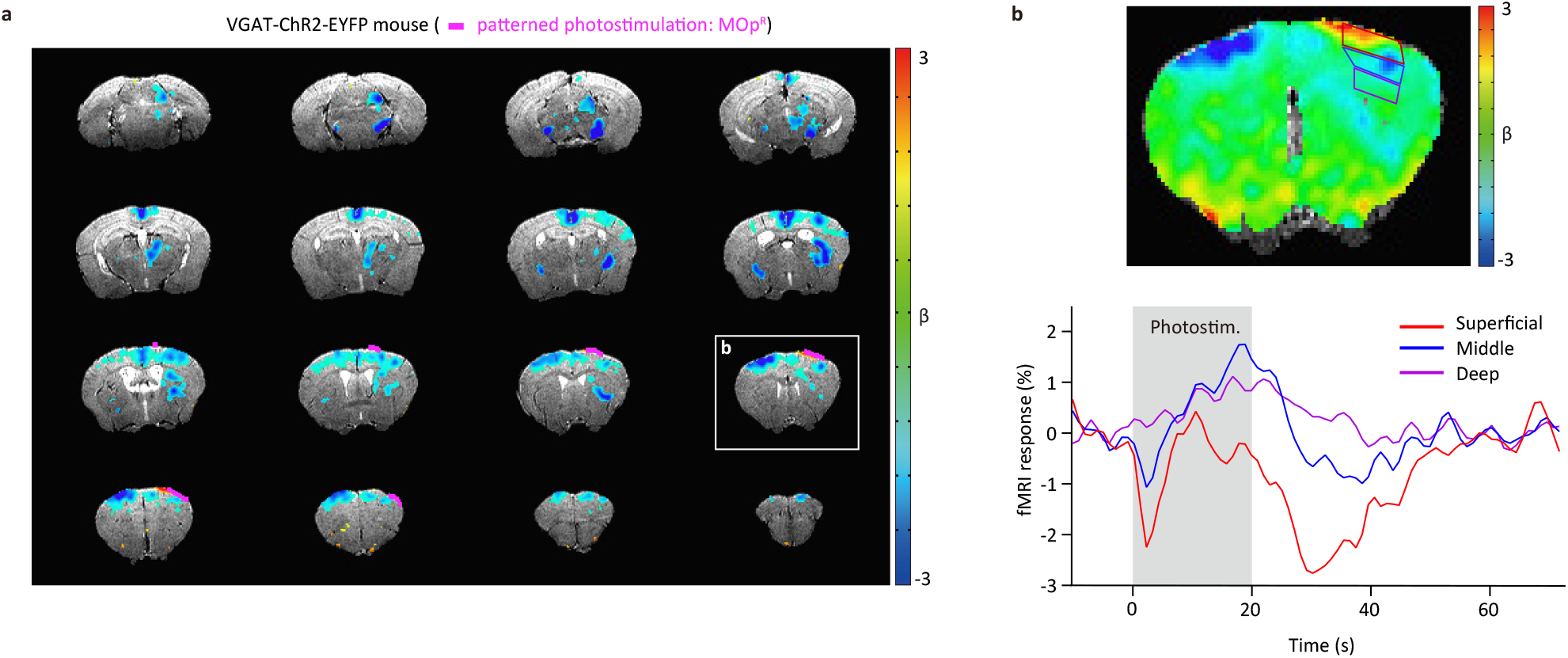
Optogenetic fMRI with patterned optogenetic stimulation of inhibitory neurons in a transgenic VGAT-ChR2-EYFP mouse. **a**, The CBV-weighted fMRI activation map in response to the optogenetic stimulus at the right MOp (20 s, 5.5 mW/mm^2^). The negative beta value represents a decrease in CBV by optogenetic stimulation. The distribution of activated voxels was highly similar to that acquired from the optogenetic excitation of excitatory neurons (Thy1-ChR2-EYFP mice) but with the opposite polarity. The right MOp, the photostimulated ROI, is marked by a purple line. **b**, To better examine the CBV response in the stimulated site, an unthresholded beta map of a selected slice in (boxed slice in **a**) is shown. At the cortical surface in the stimulated site, a positive CBV response was detected (red pixels), whereas the deeper region had a negative CBV response (blue and purple borders for the middle and deep regions, respectively). Thus, depth-dependent time traces were extracted from the 3 depth ROIs. CBV changes in the superficial and middle areas showed distinctive biphasic responses, whereas a decrease in CBV (i.e., an increase in CBV-weighted fMRI signal) was observed in the deep area. These responses can be explained by a combination of vasodilation by direct inhibitory neural activity in the stimulating site and vasoconstriction by decreased excitatory neural activity^50^. The shaded bar indicates the duration of photostimulation.

**Extended Data Fig. 10.**
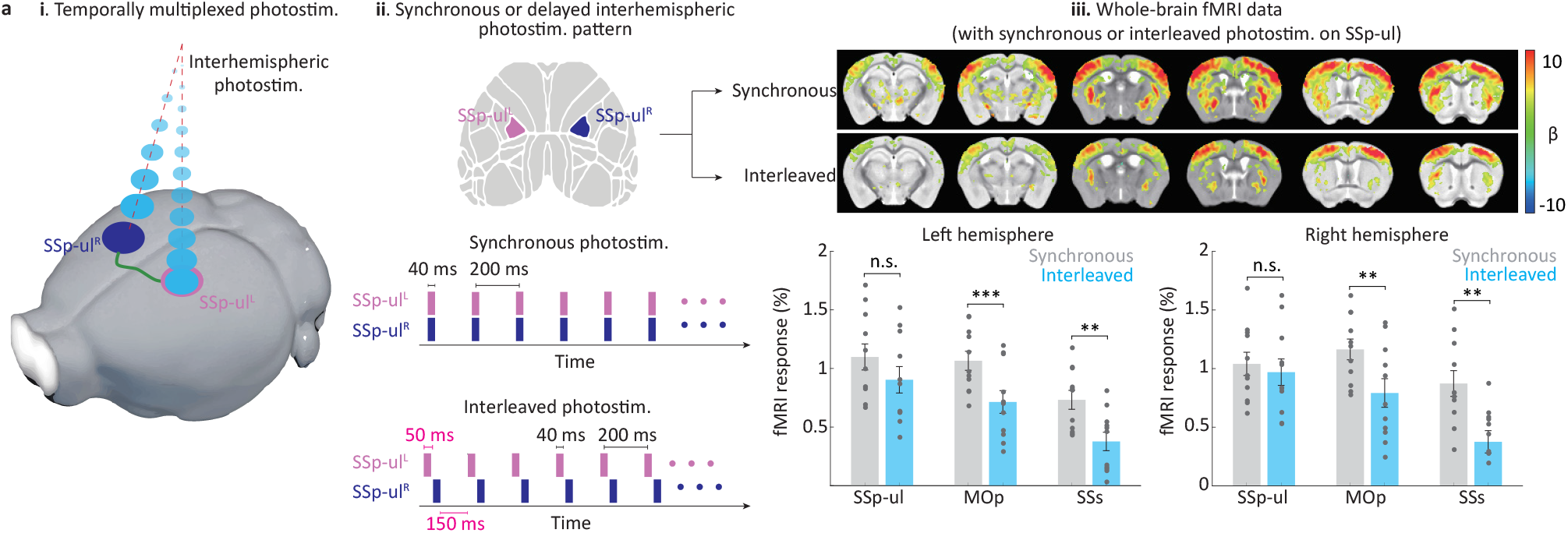
Investigation of interhemispheric neural interactions induced by precise spatiotemporally controlled optogenetic stimuli to bilateral cortical regions. (i) Interregional brain coupling of left and right forelimb activity was investigated by delivering simultaneous or interleaved photostimulation. (ii) Two stimulation designs with 5-Hz and 40-ms pulses were performed on the SSp-ul: simultaneous bilateral stimulation and interleaved bilateral stimulation with a 50-ms delay (left -> right SSp-ul). (iii) fMRI activation maps and regional responses to bilateral optogenetic stimulation. At each condition, 11 trials were repeated, and their individual responses are reported in the bottom panels. Compared to synchronous stimulation, interleaved stimulation significantly reduced the fMRI response in the MOp and SSs, while no significant difference was found in the bilateral SSp-ul. The Wilcoxon signed-rank test was conducted for statistical analysis; ***, p<0.001; **, p<0.01; *, p<0.05.

## Methods

### Optics system

The optogenetic light source emitted from the fiber-coupled laser (GP-450-100-20, RPMC) was collimated to be a 4-mm diameter circular shape. We expanded the beam diameter 3-fold with a beam expander (GBE03-A, Thorlabs). The expanded beam was directed to the center of the active DMD window (DLP9000, Texas instrument) by a pair of mirrors. The shape of a circular beam was patterned by modulation of the programmable micromirror pixels (2560×1900 pixels, refresh rate up to 9.5 kHz). We coupled the patterned beam to the fiber bundle (100,000 cores, 1.6 mm outer diameter, 3 meters long; FIGH 100-1500N, Fujikura) by passing through a tube lens (TTL200A, Thorlabs), a dichroic mirror (T470lpxr, Chroma), and an objective lens (10x, NA0.25, PLN10x, Olympus; **Fig. 1b** and **Extended Data Fig. 1a**). A filter wheel with a longpass filter (FF01-496/LP, Thorlabs) was placed in front of the sCMOS camera to allow for switching between fluorescence and reflection modes. The optical imaging signals emitted from the fiber bundle were imaged by a sCMOS camera (2048×2048 pixels, Zyla 4.2, Andor) through an objective lens and a tube lens (**Extended Data Fig. 1b**). A 530 nm LED (M530L4, Thorlabs) for reflectance and OIS imaging was coupled to a multimode fiber-optic patch cord (MFP 600/660/2000-0.22 10 m SMA-ZF1.25, Doric Lenses) and modulated by an LED driver (DC4104, Thorlabs) with the external modulation inputs from outside the MR room. Input patterns transmitted to a DMD digital controller (DLPC900, Texas Instruments) via DMD control software (DLP-LightCrafter, Texas Instruments) shaped the circular beam to a series of patterned optogenetic stimuli. The photostimulation frequency was modulated by a motorized shutter (SH1, Thorlabs) or a function generator. All optic components were fixed by screws on an optic breadboard (60×90 cm, MB6090, Thorlabs). Additionally, the optical breadboard was tightly fixed approximately 2.5 meters away from the MR bore. The timing trigger and frequency modulation control of the laser, shutter, DMD, forepaw electric stimulation, and camera were controlled by the TTL control device (Master9, AMPI), which was synchronized with MRI data collection. An achromatic projection lens (ACH 6.25 × 7.5 VIS 0 TS, Edmund optics), and the distal tip of a fiber bundle was fixed on the home-built projection module described in the main section (**Fig. 1c** and **Extended Data Fig. 1c, d**). The position of the projection lens was precisely adjusted to determine the size and position of the FOV and fixed after focusing on the reflectance image of the pial blood vessel of the mouse brain under 530 nm epi-illumination.

### Optics design and validation of the pattern projection

For precise projection of a photostimulation pattern over the cortical FOV, optics from the distal tip of the fiber bundle to the focal plane of the mouse brain were designed by sequential mode in Zemax. Briefly, the diameter of the circular projection FOV was designed to be 14 mm, which corresponds to a 10-fold magnification for the diameter of the fiber bundle. The depth of field was 2.76 mm, which was enough to cover the axial extension of the curved mouse cranium within our observation window. The dimensional parameters of the mouse cranium were adopted from the previous literature on Crystal Skull^1^ (LabMaker). The spatial resolution of the photostimulation pattern is limited by the multiplication of the single-core pitch (~4 μm) and the projection magnification (10X), which is approximately 40 μm. Optical validation of the photostimulation pattern was performed by placing the CMOS camera (CS895MU, Thorlabs) at the focal plane of the beam pattern projection (**Extended Data Fig. 3b, c**). We confirmed that both the FOV of the bundle fiber and the DMD active area covered the entire cortical window. A full-field spot diagram on a curved mouse cranium was obtained by Zemax simulation. The power density of the photostimulation pattern was measured by placing the power meter sensor (PM121D). The beam irradiance used in this study was up to 32 mW/mm^2^.

### Availability of the parts (cost) and stability of the optical setup

The optical setup is composed of an optogenetic laser source (~$15k), camera (~$15k), DMD (~$10k), fiber bundle (~$5k), lenses, LED sources for OIS imaging, and optical mounts. All parts except the projection module are commercially available from the vendors. The customized parts for the projection module can be fabricated in a local machine shop. The system, excluding the MRI machine, can be built for approximately $60k in total. No noticeable degradation in optomechanical and electrical parts was observed for ~2 years in the MR room. Occasional minor adjustment on optical alignments was required for 3-6 month intervals.

### Monte Carlo simulation

Simulations for beam profile visualization and tissue temperature change prediction were modified from previously reported MATLAB codes^2^. For the volumetric visualization of patterned illumination, the circular beam profile of a single core of the fiber bundle was convoluted with a star-shaped pattern^3^. For the prediction of the tissue temperature changes, we conservatively set the irradiance for the patterned stimulation to 5.0 mW/mm^2^, which is higher than the irradiance we used in this study. However, for spot illumination, we applied a slightly lower light dose than that used in the rodent ofMRI protocol^4^.

### Animal subjects

We used Thy1-ChR2-EYFP mice, which mostly express ChR2 within layer 5 pyramidal neurons (B6. Cg-Tg(Thy1-COP4/EYFP)9Gfng/J; stock #007615; Jackson Laboratory, Bar Harbor, ME; 18-30 g, 9 males and 10 females, aged 9-15 weeks) and a VGAT-ChR2-EYFP mouse (B6. Cg-Tg(Slc32a1-COP4*H134R/EYFP)8Gfng/J; stock #014548; Jackson Laboratory; 30 g, female, 32 weeks). The transgenic mice were bred in-house after the breeding pairs were initially obtained from Jackson Laboratory. Additionally, a wild-type C57BL/6 mouse (Orient Bio, South Korea; 29 g, male, 12 weeks) was used for the fMRI experiment with a viral injection (**Extended Data Fig. 3b**). Mice were housed under a 12/12-hour dark-light cycle, while food and water were provided *ad libitum.* All procedures were approved by the Institutional Animal Care and Use Committee of Sungkyunkwan University in accordance with standards for humane animal care from the Animal Welfare Act and the National Institutes of Health Guide for the Care and Use of Laboratory Animals.

### Animal preparation

For thinned-skull cranial window preparation, 4% ISO was delivered to mice for anesthesia induction and reduced to 2% was delivered for anesthesia maintenance during surgery. Mice were then fixed to a stereotaxic frame (Narishige, Tokyo, Japan), and their scalps were incised to expose the entire dorsal skull. After removing the thin periosteum, the skulls were thinned with a handheld drill, and then a thin layer of cyanoacrylate glue (Loctite 401, Henkel, Düsseldorf, Germany) was applied to the entire skull surface to achieve optical clarity. We surrounded the skull surface with a wall made of dental cement with a height of ~1 mm and a diameter of ~12 mm, then filled the space between the wall and the skull surface with an agarose solution. Meloxicam (1 mg/kg) was administered subcutaneously to reduce pain and inflammation. The mice were kept in cages for 2 days for until they recovered from the experiment. For the circuit-specific optogenetic fMRI, we injected rAAV2-retro-hSyn-hChR2-EYFP (a gift from Karl Deisseroth; Addgene viral prep # 26973-AAVrg) into the left inferior colliculus (AP −5.3mm, ML +1.0mm from the bregma), 200 nl each at a depth of 0.7 mm and 1.4 mm, to a wild-type C57BL/6 mouse. The viral injection was performed 4 weeks before the experiment.

For MRI experimental preparation, mice were initially kept in a chamber with 4% ISO for ~4 min for anesthesia induction. In the case of ketamine/xylazine anesthesia^5^, a ketamine (100 mg/kg) and xylazine (10 mg/kg) mixture was first injected intraperitoneally (IP) and maintained by intravenous (IV) continuous infusion of ketamine (35-45 mg/kg/hr) and xylazine (1.75-2.25 mg/kg/hr) into the tail vein during experiments. For the ISO experimental group, 1% ISO was maintained throughout the preparation and experiment. The skull surface was cleaned carefully to remove dust, which can prevent light stimulation. The dental resin well was filled with a 0.5% agarose solution in D2O to minimize MRI artifacts, and three or four fluorescence tubes (small pieces of a PE-10 tube filled with rhodamine solution in H2O) were placed inside the agarose gel as a reference for coregistration between MRI and optical images (**Fig. 2a, b**). Note that MRI and fluorescence imaging can detect the fluorescence tubes, whereas neither can detect the agarose gel. The well was covered by a cover glass, and the border was thoroughly sealed using biocompatible silicone adhesive (Kwik-Sil, World Precision Instrument, Sarasota, FL) to prevent the evaporation of agarose solution. The size of the cranial window did not exceed a diameter of 15 mm so that it fits within the RF coil (inner diameter of 15 mm) (**Fig. 2a**). A 31G needle connected to a 25G Y-connector (SCY25, Instech Laboratories, Plymouth Meeting, PA) was inserted into the tail vein to deliver anesthetics and the contrast agent (monocrystalline iron oxide nanoparticles, MION; Feraheme, Waltham, MA) for CBV-weighted fMRI.

### Physiology monitoring

Each mouse was positioned in a customized mouse cradle, and the head was fixed with ear and bite bars to reduce motion artifacts. To maintain the animal’s condition, 1) a respiration-sensitive sensor was attached to the belly, 2) a temperature sensor was inserted into the rectum, and 3) a pulse oximeter was attached to the tail (**Fig. 1b**); measurements from all of these devices were monitored with an animal monitoring system (Model 1030, Small Animal Instruments, Stony Brook, NY). During the experiments, the animal’s body temperature was kept at ~37°C using a cradle-embedded hot water tube and a water heating pad, and oxygen-rich air (a mixture of oxygen and air at a 1:4 ratio) was ventilated continuously through a nose cone at a rate of 0.4 mL/min using a small animal ventilator (TOPO, Kent Scientific Corporation, Torrington, CT). For CBV-weighted fMRI, MION (20 mg/kg) was injected intravenously. A stimulation-evoked blood volume increase induced an increase in superparamagnetic iron oxides within the voxel and consequently decreased fMRI signals. This CBV-weighted fMRI approach enhances the sensitivity and specificity of fMRI responses.

### MRI data acquisition

MRI data were acquired on a 9.4 T/30 cm MR scanner (Bruker Biospec, Billerica, MA, USA) with an actively shielded 12-cm gradient operating with a maximum strength of 66 G/cm and a rise time of 141 μs. An 86-mm volume coil for RF transmission and a single-loop surface coil with an inner diameter of 15 mm and an outer diameter of 20 mm (Bruker Biospec, Billerica, MA, USA) were used. First, a scout image was acquired to localize the position of the fluorescence tubes. Then, a 3D T2-weighted anatomical image was obtained using rapid acquisition with refocused echoes (RARE) with the following parameters: FOV = 16 (readout, L-R) × 10 (V-D) × 10 (A-P) mm^2^ (including the position of fluorescence tubes); spatial resolution = 0.1×0.1×0.1 mm^3^; repetition time (TR)/echo time (TE) = 1500/36 ms; RARE factor = 16; and total scan time = 16 min and 30 s. CBV-weighted functional images were acquired using either the 2D or 3D single-shot gradient-echo echo-planar imaging (EPI) sequence with the following parameters: For 2D EPI, FOV = 16 (readout, L-R) × 8 (V-D) mm^2^; 18 contiguous coronal slices with a thickness of 0.5 mm; in-plane resolution = 0.167 × 0.167 mm^2^; TR/TE = 1000/8.35 ms; flip angle (FA) = 47°; and scan time for one fMRI run ≤ 440 s. For 3D EPI, FOV = 16 (readout, (L-R) × 12 (A-P) × 8 (V-D) mm^2^; spatial resolution = 0.2 mm^3^; TR/TE = 50/8 ms (temporal resolution of 2 s); flip angle (FA) = 12°; and scan time for one fMRI run ≤ 580 s. In the experiments of whole-brain mapping with stimulation of 9 atlas-based ROIs (**Fig. 3**, **Fig. 4**, and **Fig. 5**), the fMRI data were acquired with 3D EPI, while otherwise with 2D EPI. For EPI, local shimming was performed based on the B_0_ map to minimize field inhomogeneity within the elliptical shim volume inside the brain. Saturation of the ventral slab outside the FOV was performed to alleviate the aliasing effects along the phase-encoding direction.

### Generation of subject-specific stimulus patterns with coregistration of optics and MRI

The active windows of the DMD and sCMOS sensor areas were first aligned and coregistered before the MR experiment via a point-based registration algorithm. First, an arbitrarily shaped polygon pattern (we used a star shape) was displayed on the DMD window. Second, the pattern that reflected the bundle fiber end was imaged by sCMOS. Finally, multiple corners of the polygon pattern image were selected and transformed in MATLAB to be the same as the input pattern. After this process, we successfully obtained the transformation matrix between the sCMOS image and the DMD input pattern (M3 matrix, **Fig. 2d**). Matrix M3, corresponding to the optical alignment on the optic breadboard in the MR room, was not affected by magnetic fields for at least 6 months. The subject-specific optogenetic stimulus pattern was generated by the following procedure. First, a T2-weighted anatomical MR image was acquired, which also contains the location of reference tubes. A customized mouse brain template^6^ that shares the same space with the Allen Mouse Brain Atlas (R^Allen^) was coregistered onto the anatomical image (ANTs antsRegistrationSyN, R^MR3D^) to obtain the M_1_ transformation matrix. Then, only the most superficial voxels of 3D MRI (R^MR3D^) were projected (M2Dprj) onto a dorsal optical plane (i.e., y-axis in MRI). The resulting projected image (on the native MRI space) (R^MR2D^) was aligned with the fluorescence image with affine transformation (MATLAB fitgeotrans) using the locations of fluorescence tubes as reference points (R^Fluor^) to obtain the M2 transformation matrix. Finally, DMD input patterns for the cortical regional photostimulation were generated by multiplying the Allen Atlas data with a combination of the multiple transformation matrices (M_1_, M_2_, M_2Dprj_, and M_3_) (**Fig. 2e**). For Allen Atlas-based stimulation, 56 dorsal cortical regions in both hemispheres were parcellated based on the definition of the nine ROIs chosen for optogenetic stimulation (**Fig. 3f**). Alternatively, the optogenetic stimulus target can be determined directly from optical or MRI with M2 and M3 matrices. For example, forepaw stimulation-evoked intrinsic optical images were used to identify forelimb areas and determine optogenetic stimulation patterns (**Extended Data Fig. 3a**), while fluorescence optical imaging was used to localize and target the cortical expression of ChR2 by retrograde viral injection into the IC (**Extended Data Fig. 3b**).

### Stimulus paradigm

A series of optogenetic stimulations of 9 cortical regions was performed sequentially with stimulation and poststimulus durations of 10 s and 50 s, respectively, (**Fig. 3, 4, 5**) during a single fMRI run or separate runs. The order of the optogenetically activated regions was fixed in one animal for straightforward averaging but was changed for different mice. Except for dose-dependent fMRI studies (**Fig. 3**), optogenetic stimulation parameters were fixed with a power density of 3.6 mW·mm^-2^ for further experiments in **Fig. 4** and **Fig. 5**. Moreover, we used a stimulation frequency of 20 Hz and a pulse duration of 10 ms (except for **Extended Data Fig. 10**; 5 Hz and 40 ms). For 20-s forepaw somatosensory activation studies (**Extended Data Fig. 3a**), the stimulation parameters were a current of 0.5 mA, frequency of 4 Hz, and pulse duration of 0.5 ms.

### Functional MRI preprocessing and activation maps

All procedures for image processing and data analysis were performed using the following software packages: Analysis of Functional NeuroImages^7^ (AFNI), FMRIB Software Library^8^ (FSL), Advanced Normalization Tools^9^ (ANTs), and MATLAB (MathWorks, Natick). Repeated fMRI runs were averaged in each subject. The EPI dataset from every animal was first preprocessed with slice timing correction and motion correction and were coregistered to a T2-weighted anatomical image with rigid-body transformation (FSL flirt). Individual datasets were normalized to the customized brain template^6^ (on the same coordinate with the Allen atlas) using the transformation matrix generated in the previous section (ANTs antsApplyTransforms).

We defined the data-driven hemodynamic response function (HRF) for the generation of fMRI activation maps. First, in the dose-dependent experiments (**Fig. 3**), we extracted time courses from atlas-based ROIs (65 ROIs x 2 hemispheres x 9 stimuli x 5 animals; **Fig. 3h**) in stimulus-free fMRI runs and computed the mean % change of each ROI. Based on this null distribution, we determined the threshold to be −1.28% (corresponding to a Z score of 3.09; p<0.001 for the one-tailed z test) for the presence of a response. Then, time traces with a mean signal change larger than −1.28% were selected from the fMRI runs with optogenetic stimulation (69, 107, and 291 traces for 1.2, 2,4, and 3.6 mW/mm^2^, respectively) and averaged. The averaged response was then fitted by a double gamma variate function to determine the HRF. The same procedure was also conducted on the K/X (n = 8 mice) and ISO datasets (n = 9 mice), resulting in anesthesia-dependent HRF (**Fig. 5d**). Then, the individual activation maps were generated after spatial smoothing with a Gaussian kernel (full-width at half maximum (FWHM) of 0.156 mm or 0.2 mm: one voxel size of 2D and 3D EPI), using general linear model (GLM) analysis conducted with a design matrix consisting of stimulation periods of HRF with linear detrending (AFNI 3dDeconvolve).

The group-averaged activation maps were generated in two ways. For the dosedependent activation map (n = 5 mice; **Fig 3i**), we made the maps by applying GLM to the concatenated individual time series datasets to capture the fixed effect of the stimulation dose (AFNI 3dDeconvolve). For the activation maps of the K/X and ISO groups (**Fig. 4b**, **Extended Data Fig. 4**, and **Extended Data Fig. 6**), we conducted a voxel-by-voxel t-test across individual activation maps in each group with cluster-extent-based multiple comparison corrections (AFNI 3dttest++ with Clustsim).

### Quantitative analysis for effective and structural connectivity

For the structural connectivity data, we imported 9 representative experiments from the Allen Mouse Connectivity Atlas (https://connectivity.brain-map.org; Experiment # 127084296, 112952510, 112229814, 112951804, 112881858, 307593747, 657334568, 100141599, and 100140949), presented as projection volumes in each Allen Mouse Brain Atlas-based structure. The connections with a projection volume smaller than 0.008 mm^3^ (one voxel volume) were excluded. The projection volume was recalculated to conform with the fMRI ROI definition (i.e., summation of data in the regions within the ROI). We presented ROI-level fMRI data in two ways. For the comparison of EC and SC (**Fig. 4** and **Extended Data Fig. 7**), we computed EC as a sum of voxel-by-voxel responses in each ROI divided by a sum of voxel-by-voxel responses of the stimulated ROI, while SC was computed as a projection volume in each ROI divided by the projection volume of the source ROI. Otherwise, the fMRI data were presented in a typical manner: the mean of voxel-by-voxel responses in each ROI.

### Statistics

Statistical analyses in this paper were performed using MATLAB 2019b or R version 4.1.2 and RStudio version 2021.9.1.372. All quantitative values are presented as the mean and standard error of the mean. For the calculation of the correlation between log EC and SC (MATLAB *corr*: **Fig. 4f-h** and **Extended Data Fig. 7c-e**), zero and negative values in the original data (on a linear scale) were assigned to −10 (on a log scale), the lowest value in any dataset. PERMANOVA (**Fig. 5f**) was performed using the *vegan* version 2.5-7 package on the Euclidean distance matrix (999 permutations). PCA was performed using MATLAB *pca* with individually normalized response matrices (**Fig. 5g, h**). For the PLS-DA, the package *ropls* version 1.26.4 was used on the unscaled stimulation-induced response matrix (**Fig. 5i** and **Extended Data Fig. 8**). The PLS-DA model with two components (**Extended Data Fig. 8a**) was validated using fivefold cross-validation (999 permutations). The first two components were used to calculate VIP values for each stimulation-induced response matrix. PLS-DA score plots were visualized using the package *ggplot2* version 3.3.5. The resulting VIP value data matrix was visualized as a heatmap using the package *corrplot* version 0.92 (**Extended Data Fig. 8b**). Throughout this paper, t-tests and Mann–Whitney U tests with FDR adjustment were performed using the *stats* package in base R.

## Data availability

All relevant data are available from the corresponding authors upon request. A reporting summary for this article is available as a Supplementary Information file.

## Code availability

The software codes are available online at http://github.com/cnir-sgk/DMD-fMRI-v1.0.

## Supplementary Information

**Supplementary Figure 1.**
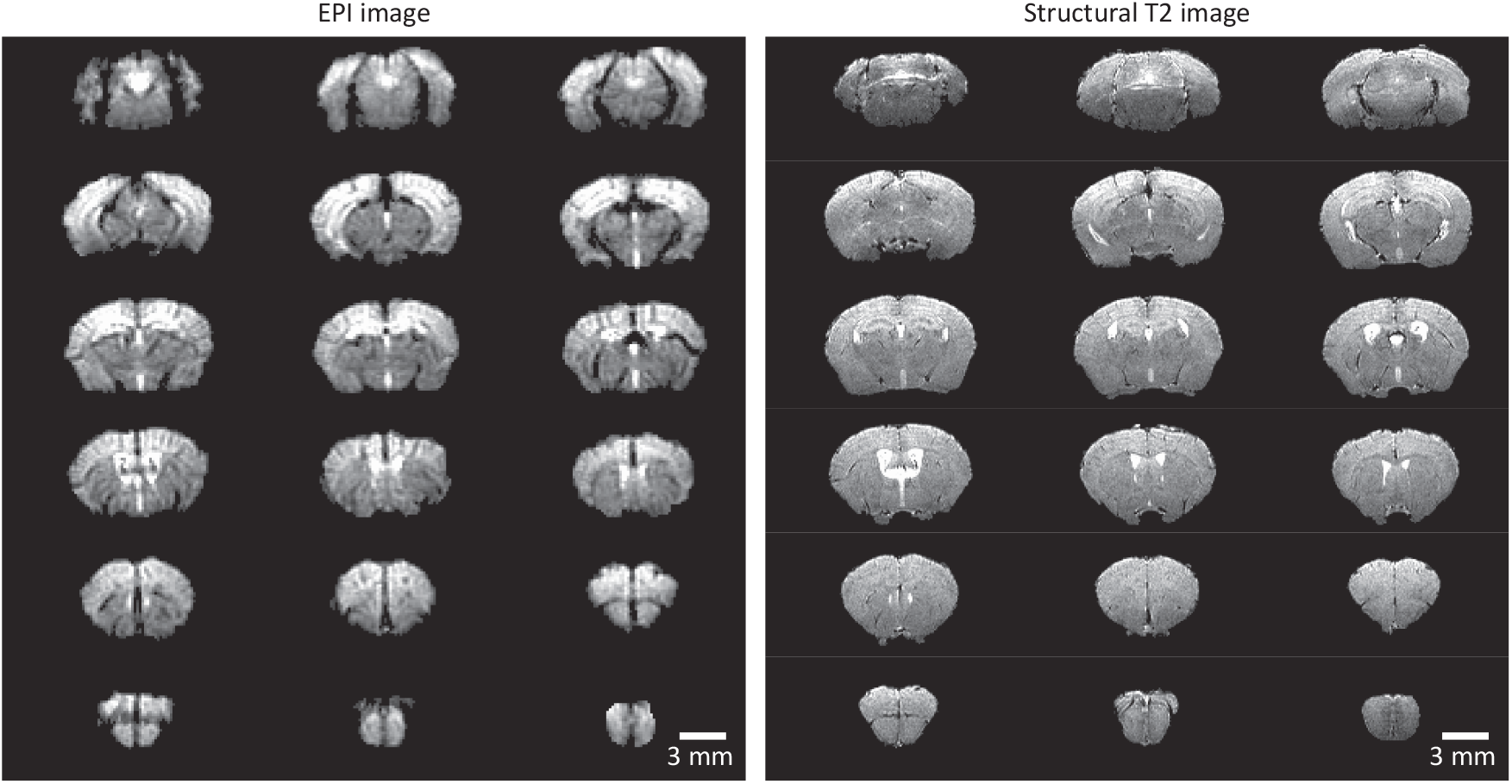
Whole-brain MR functional (gradient-echo EPI) and structural T2-weighted images (RARE) of one representative thinned-skull mouse. Neither MR artifacts nor image distortions were observed in a gradient-echo EPI image since D2O-based agarose gel covering the cranial window minimizes the susceptibility effect and enhances shimming. Both types of images obtained after MION injection clearly show the vasculature of the brain.

**Supplementary Video 1 | Schematic video of the experiment.** The DMD input pattern sequence, experimental scheme, color video (taken outside the MR and synced later) and its corresponding fMRI activation 3D maps are graphically presented.

**Supplementary Video 2 | OIS imaging through the fiber bundle under fMRI acquisition.** The video was reconstructed from a series of reflectance images acquired through the fiber bundle during fMRI recording under electrical forepaw stimulation.

